# On the road to losing connectivity: Fecal samples provide genome-wide insights into anthropogenic impacts on two large herbivore species in central India

**DOI:** 10.1101/2023.10.26.564122

**Authors:** Abhinav Tyagi, Nidhi Yadav, Awadhesh Pandit, Uma Ramakrishnan

**Affiliations:** National Centre for Biological Sciences, Tata Institute of Fundamental Research, Bangalore, India; SASTRA Deemed to be University, Thanjavur, India

**Keywords:** ddRAD, Non-invasive samples, Gaur, Sambar, Wildlife genomics, Wildlife connectivity

## Abstract

Humans have impacted most of the planet, and the ensuing fragmentation results in small, isolated habitat patches posing a risk of genetic diversity loss, inbreeding and genetic load. Understanding how natural and anthropogenic landscape features affect gene flow among habitat patches is critical for maintaining connectivity. Genome-wide data is required to comprehend the impacts of recent fragmentation, which can be challenging when only non-invasive samples are available. Here, we build upon advancements in conservation genomics to address connectivity of two large herbivores, gaur (*Bos gaurus*) and sambar (*Rusa unicolor*) in central India. Given their habitat associations, we expected these species to respond similarly to habitat fragmentation. We used fecal-DNA and methylation-based host-DNA enrichment with modified ddRAD protocol to generate genome-wide single nucleotide polymorphism (SNP) data for 124 Gaur and 99 Sambar individuals. Our findings reveal that gaur populations in central India are fragmented, displaying high genetic differentiation, with drift significantly affecting small populations like Umred-Karhandala Wildlife Sanctuary. Although Sambar shows low genetic structure, another small population, Bor Tiger Reserve is genetically differentiated. Our results suggest that although landcover change and roads restrict animal movement, the extent of this impact varies across the two species. We show that different species, respond differently to landscape features, even with similar habitat associations. We highlight small and isolated populations requiring urgent conservation intervention. Such multi-species approaches enhance our understanding of cross-species connectivity patterns. We suggest shifting from single-species to multi-species holistic conservation approach in rapidly developing landscapes to better manage co-occurring endangered species.

## 1. Introduction

The ongoing wildlife crisis leading to local and global species extinctions is primarily driven by anthropogenic impacts such as habitat loss and fragmentation (Dirzo et al., 2014; Young et al., 2016). In particular, large mammals bear the brunt of vulnerability due to their life history traits, including large home ranges, migratory behaviour, slow reproductive rates, and sparse population densities (Cardillo et al., 2004). While large carnivores have received frequent spotlight as threatened keystone species, herbivores are often neglected. However, large herbivores play a critical role in shaping ecosystem structure and function by controlling wildfires, cycling nutrients, and maintaining vegetation dynamics (le Roux et al., 2020; Rouet-Leduc et al., 2021; Sandom et al., 2014). Large herbivores also make habitats suitable for carnivores and scavengers as they are the primary food source (Ripple et al., 2015).

Unfortunately, despite their ecological significance, most large herbivore species are experiencing declines in abundance and distribution due to anthropogenic pressures such as land-use change, hunting, competition, and disease transmission (Bluhm et al., 2023; Craigie et al., 2010; Ripple et al., 2015). Land-use change, such as habitat loss, fragmentation, and deforestation, restricts movement and disrupts gene flow (Crego et al., 2021; Ripple et al., 2015). The development and presence of other anthropogenic landscape features like road and railway networks can lead to increase in habitat fragmentation and reduces landscape permeability (Bennett, 2017; Hepenstrick et al., 2012). There have been several incidences of local extinction and extirpation of large herbivores from fragmented habitat patches (Sankar et al., 2013). Therefore, maintaining gene flow and functional connectivity among these fragmented habitat patches is critical for long-term species persistence (Thatte et al., 2018).

Most connectivity studies, especially in South Asia, have focused on a single species, mostly charismatic carnivores (Thatte et al., 2021). Connectivity is often species-specific and depends on species traits and ecology like body size, dispersal ability and habitat requirement (McClure et al., 2016; Thatte et al., 2020). This is also relevant for large herbivores as they have specific habitat associations and requirements, and can exhibit high association with the migration routes or corridors that might not necessarily support the movement of other species (Sawyer et al., 2019). Therefore, a multi-species and comparative approach is essential to comprehensively understand how connectivity varies across species. This comparative analysis, especially among species with similar habitat associations, sheds light on the role of species ecology, beyond just habitat, in shaping connectivity (Brodie et al., 2015; Thatte et al., 2020, 2021).

Landscape genetics integrates methods from population genetics and landscape ecology to examine the impact of landscape features on gene flow and genetic diversity (Storfer et al., 2007). These methods are instrumental for identifying barriers to animal movement and have been applied to a diverse range of species (Dudaniec et al., 2016; Fedorca et al., 2020; Schoen et al., 2022; Thatte et al., 2020; Tyagi et al., 2022). The use of molecular markers and landscape genetic approaches aid in quantifying regional patterns of genetic structure, providing insights into how different landscape features influence connectivity (Epps & Keyghobadi, 2015; Holderegger & Wagner, 2008; Tyagi et al., 2022). In many places including India, non-invasive samples, such as fecal samples, are the only feasible means of studying large herbivores in their natural habitats. However, the DNA quality of these samples is often compromised, as fecal DNA extracts tend to contain exogenous DNA and PCR inhibitors (Khan & Tyagi, 2021; Kohn & Wayne, 1997; Putman, 1984; Schrader et al., 2012).

Recent developments in conservation genomics enable the use of DNA obtained from poor-quality non-invasive samples to generate genome-wide data using next-generation sequencing (NGS) (Chiou & Bergey, 2018; Hayward et al., 2022; Natesh et al., 2019; Orkin et al., 2020; Tyagi et al., 2022). The use of single nucleotide polymorphism (SNP) markers has led to better estimates of genetic diversity and population structure (Bohling et al., 2019; Lemopoulos et al., 2019; Ling et al., 2020) and inference of landscape features on animal movement (Tyagi et al., 2022). The use of a higher number of SNP markers provides more power in comparison to increasing the number of samples or individuals in quantifying fine-scale genetic variation and functional connectivity, as evidenced from both simulations and empirical data (Landguth et al., 2012; Tyagi et al., 2022).

Further, double-digest restriction-associated DNA (ddRAD) sequencing has emerged as a promising approach for generating genome-wide data for non-model species. Despite the promising potential, there is still a need to develop library preparation methods that complement the use of poor-quality, low-concentration input DNA and enable the generation of a comparable amounts of data per sample. Most available protocols are not designed to deal with the specific issues related to analysing samples with poor-quality and varied DNA concentrations (Peterson et al., 2012; Peterson et al., 2014). To address this, we developed a modified ddRAD sequencing protocol that allows us to generate genome-wide data, suitable for poor-quality DNA samples and minimizes the impact of input host DNA concentration on the data generated.

In order to address gaps in our understanding of how fragmentation impacts large herbivores, we utilized genome-wide SNPs obtained from fecal samples to investigate connectivity patterns of two large vulnerable herbivore species, gaur (*Bos gaurus*) and sambar (*Rusa unicolor*), in central India. Our main objectives were to (a) determine the genetic diversity and population structure of these species, (b) assess the impact of landscape heterogeneity on gene flow, and (c) compare the response of two species with similar habitat dependency, but different body size and dispersal-related functional traits, to habitat fragmentation and different landscape features. We predict that both species may reveal similar patterns of connectivity as they share similar habitat associations. Additionally, we aimed to identify priority areas for restoring connectivity for these species. This study represents the first genome-wide investigation of landscape-level connectivity for both species and underscores the importance of studying genetic variation and connectivity among different species.

## 2. Materials and Methods

### 2.1. Study area

The central Indian landscape (CIL) is a well-studied landscape in terms of connectivity studies in India, but the focus of these studies has largely been on carnivores, specifically, tigers (Schoen et al., 2022; Thatte et al., 2021). It is a priority tiger conservation landscape and is also inhabited by various other endangered wild species (Jhala et al., 2008). Our study landscape was spread across two central Indian states— Madhya Pradesh and Maharashtra. The forest is primarily tropical dry and moist deciduous, harbouring rich biodiversity (Champion & Seth, 1968). Central India has around 33% of the area under forest cover, which consists of multiple protected areas (PAs), reserved and territorial forests (Dutta et al., 2018). A varied and heterogeneous landscape matrix with multiple land-use types, such as agricultural fields and human built-ups, surrounds these protected areas.

Central India, like other areas of conservation concern, faces threats including habitat fragmentation, degradation, and loss of connectivity due to rapid transformations driven by human activities such as transportation network expansion, mining projects, and development (Crooks et al., 2011; Habib et al., 2016; Ibisch et al., 2016; Schoen et al., 2022; Venter et al., 2016). Major state and national highways bisecting the landscape further jeopardizes wildlife connectivity and integrity (Thatte et al., 2018, 2020).

### 2.2. Study species and sample collection

Gaur (*Bos gaurus*) and sambar (*Rusa unicolor*) are endemic to South and Southeast Asia and rank among the top prey species for large carnivores like tigers (Andheria et al., 2007). Gaur is the largest wild cattle species and one of the few mega-herbivore species that inhabit the Indian subcontinent. Although Gaur was once found throughout the forested areas of India, its current distribution is restricted to fragmented habitat pockets, primarily in the Western Ghats, the central Indian highlands, and Northeast India (Musser & Koop-man, 1994).

On the other hand, Sambar is a meso-herbivore distributed across a wide range of habitats, from tropical rainforests to subtropical mixed forests in South and Southeast Asia. In India, it can be found from the foothills of the Himalayas in the north to the Western Ghats in the south and the Peninsular and north-eastern regions (Timmins et al., 2015). Although sambar exhibits a wide range, the population is very patchy, especially in central India (Kushwaha et al., 2004; Timmins et al., 2015).

Both species have similar habitat association and prefer habitats with high tree and shrub densities, and generally avoid anthropogenic disturbances (Sankar et al., 2013; Singh et al., 2021). Both species face threats like habitat fragmentation, habitat loss, illegal poaching and other anthropogenic impacts responsible for population decline and local extirpation. Gaur has experienced several local extinctions in central India, including from Bor Tiger Reserve, Bandhavgarh Tiger Reserve, and Kengar Valley National Park (Sankar et al., 2013). While there are no published reports of local extinctions for Sambar, the lack of monitoring and attention towards large herbivores raises concerns about its status in many areas.

To assess the genetic connectivity of the two species, we collected fecal samples during two field seasons (October 2017 to February 2018 and December 2020 to March 2021) from inside and outside six protected areas: 1) Kanha Tiger Reserve (KTR), 2) Pench Tiger Reserve (PTR), 3) Nagzira-Nawagaon Tiger Reserve (NNTR), 4) Bor Tiger Reserve (BOR), 5) Tadoba-Andhari Tiger Reserve (TATR), and 6) Umred-Karhandla Wildlife Sanctuary (UMR). Outside protected area sampling was done in forested areas like the Kanha-Pench corridor (KPC) (Figure 1).

**Figure 1:**
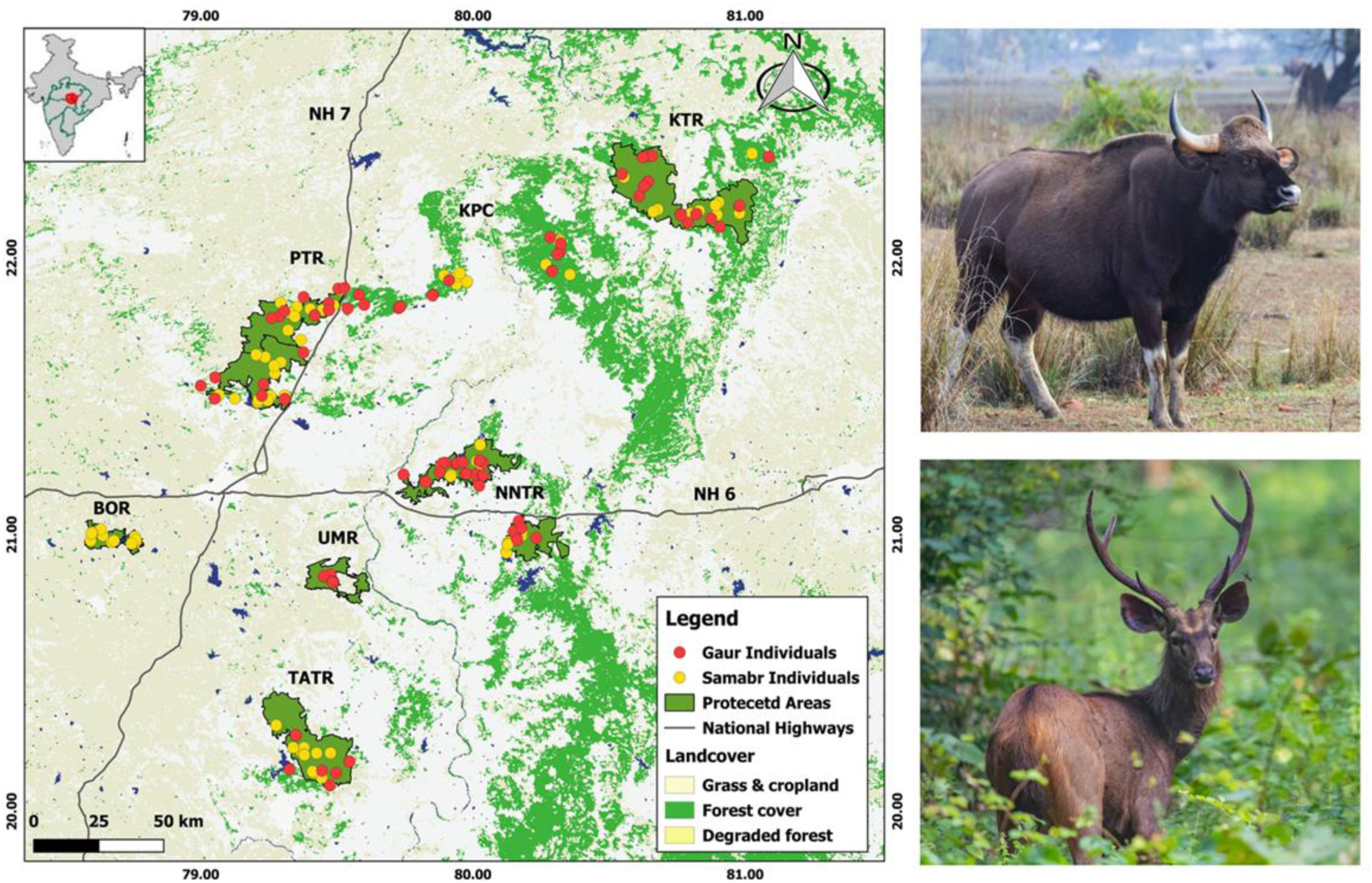
Study area with sampling sites. Red and yellow dots represent gaur (top right) and sambar (bottom right) individuals respectively. Two major national highways (NH6 & 7), bisecting the study landscape are also shown as bold black lines. Protected areas are marked in dark green colour with black boundaries. KTR – Kanha Tiger Reserve; KPC – Kanha Pench corridor; PTR – Pench Tiger Reserve; NNTR – Nagzira Nawegaon Tiger Reserve; BOR – Bor Tiger Reserve; TATR – Tadoba Andhari Tiger Reserve and UMR – Umred Karhandala Wildlife Sanctuary.

We sampled in areas with frequent use by study species to maximize the collection of fresh samples (less than 3 days old). Fresh dung (for gaur) and pellets (for sambar) were sampled using a sterilized swab and stored in Longmire’s buffer in 2 ml microcentrifuge tubes (Longmire et al., 1997). Each sample was collected in duplicate, and we targeted a large sample size to compensate for a potential loss of samples due to the low quality and quantity of DNA. To maximize the area covered and avoid recaptures of individuals, each forest beat (i.e., administrative divisions) was sampled only once, and adjoining beats were sampled simultaneously. We recorded GPS location, species identity based on morphology, and a visually estimated sample age for all samples.

### 2.3. DNA extraction, species identification and quantification

All samples were extracted using QIAamp DNA Blood Mini Kit (Qiagen) following the manufacturer protocol. Species identification was done for all the samples using species-specific primers designed in this study. The gaur (Bg_cytb1.1F *5’-CACAGCAATTGCCATAGTCC-3’* & Bg_cytb1.1R *5’-AAGGGCTAGAATTAGTAACAAGGTT-3’*) and sambar (CommF_CytB1 *5’-ATTGTAAACAACGCATTCAT-3’* & SambarR_CytB1 *5’-GGAATAGGCCTGTGATG-3’*) specific primers amplified 151 and 115 base pair sequences, respectively, of mitochondrial cytochrome b region (Table S1 and Supplementary material S1). No cross-amplification, with any of the other sympatric species with which the fecal samples could be misidentified based on morphology, was observed. Therefore, any positive amplification observed using fecal DNA corresponds to conclusive species identity (Supplementary material S1).

Total DNA concentrations were measured post species identification, using Qubit (Invitrogen). Host DNA (hDNA) was estimated using quantitative PCR (qPCR) (Morin et al., 2001). All qPCR reactions were performed in triplicates along with positive and negative controls and evaluated against the standard curve constructed using serial dilutions of known tissue DNA.

### 2.4. DNA enrichment

Host DNA enrichments were performed using the NEBNext Microbiome DNA enrichment kit (Feehery et al., 2013). The samples and MBD2-Fc magnetic beads were prepared following the procedure described in Tyagi et al., (2022). To maximize the hDNA capture, the DNA extracts were concentrated using Cenrivap DNA vacuum concentrator (Labconco). Total and hDNA were also quantified post-enrichment.

### 2.5. Library preparation and ddRAD sequencing

We prepared ddRAD libraries using the enriched fecal DNA samples. Our protocol was based on modifications of existing methods (Peterson et al., 2012; Peterson et al., 2014; White et al., 2018) and tailored to our sample types. We digested all samples with a combination of Sph1 and MluC1 restriction enzymes and performed adapter ligation, purification using AMPure XP magnetic beads (Agencourt), and indexing. To reduce PCR bias, we performed indexing PCR in four replicates and pooled all replicates before conducting dual-size selection using AMPure XP magnetic beads. We quantified and assessed the size distribution of individual sample libraries using Qubit and Tapestation (Agilent Technologies), respectively, before pooling them in equimolar concentration. The final pooled library was also quantified and checked for size distribution before sequencing on the HiSeq2500 platform using PE 2X100 V2 kit. Please see supplementary material S2 for detailed protocol, adapter, index and, modified read 1 and read 2 sequences (Table S3 and S4).

### 2.6. Quality control, SNP calling and filtering

Raw reads were first demultiplexed and then FastQC was used for quality checking (http://www.bioinformatics.babraham.ac.uk/projects/fastqc/). The reads were trimmed using TrimGalore (v.0.5.0 Babraham Bioinformatics) and mapped to respective reference genomes using the BWA mem algorithm (Li, 2013). Cow reference genome (ARS-UCD1.2; RefSeq accession: GCF_002263795.1) was used to align gaur data and red deer reference assembly (mCerEla1.1; RefSeq accession: GCF_910594005.1) was used for mapping sambar reads. We selected these reference genomes for mapping as they represented the best available reference assemblies for the species most closely related to our study species (Mackiewicz et al., 2022; Wu et al., 2018). Post mapping, read-group information was added to each BAM file using Picard tools (http://broadinstitute.github.io/ picard).

Variant calling was done using Freebayes (Garrison & Marth, 2012) and vcftools (Danecek et al., 2011) was used to filter the sites based on quality (≥10) and minor allele count (≥3). Sites supported by three or more reads were considered for identifying genotypes (Arafa et al., 2017; Chang et al., 2023; Chen et al., 2021; Shirasawa et al., 2016). Only biallelic variants were retained, and sites that were present in less than 60% of the individuals were discarded. Samples with more than 40% missing data were removed from further analysis.

### 2.7. Individual identification, isolation by distance and outlier loci detection

We generated relatedness estimates using PLINK (Purcell et al., 2007) to find recaptures and related individuals. For testing Isolation by distance (IBD), we calculated pairwise genetic distance based on the proportion of shared alleles (D_PS_=1-proportion of shared alleles) and Euclidian distance using *adegenet* package (Jombart, 2008) in R version 4.1.2. We tested for correlations between genetic and geographic distances using Mantel tests using package *adegenet* (Jombart, 2008) with randomization with 999 replicates. Genalex 6.503 (Peakall & Smouse, 2012) was used to calculate and plot a multi-locus spatial correlogram using genetic and geographic distances using 1000 bootstraps, and 1000 permutations of the data were used to evaluate significance. We also used Bayescan software (Foll & Gaggiotti, 2008) to detect and remove loci under selection, using a false discovery rate of 5%.

### 2.8. Genetic differentiation and population structure

Expected and observed heterozygosity was computed using R packages *adegenet* (Jombart, 2008) and *Hierfstat* (Goudet, 2005). We used multiple approaches to understand genetic differentiation comprehensively. We assessed each species’ genetic structure using discriminant analysis of principal components (DAPC) and STRUCTURE (Pritchard et al., 2000). DAPC (Jombart et al., 2010) was conducted using the *adegenet* package in r version 4.1.2. The number of principal components (PC) axes and discriminant functions (DFs) to be retained were determined using the optim.a.score function. We also used the program STRUCTURE (Pritchard et al., 2000) as a Bayesian approach to determine the most likely number of genetic clusters (K). STRUCTURE was run for 200,000 Markov Chain Monte Carlo (MCMC) iterations after a burn-in of 50,000 iterations, with ten replicates for K ranging from one to eight, using the correlated allele frequencies and admixture model. The best K was selected by likelihood and delta K values and methods described by Evanno et al., (2005). The results were analysed and plotted using a web-based server, CLUMPAK (Kopelman et al., 2015).

### 2.9. Landscape genetic analysis

#### 2.9.1. Landscape variables

To build resistance models in this study, we used four landscape variables: land use land cover (LULC), roads with traffic, human population density (Hpop), and density of linear features (Linden). These variables have been previously used to understand the connectivity of various species in the same landscape (Thatte et al., 2020; Tyagi et al., 2022). Each layer was reclassified and ranked in order of the resistance offered. Landcover data was reclassified into four categories: forest, scrub and degraded forest, agriculture, and built-up areas. The road layer was reclassified into seven different classes based on traffic intensity (measured in Passenger Car Units (PCU)) and road width (e.g., 1,2,4, or 6 lanes). To calculate the density of linear features, we combined roads (without traffic), railway lines, and irrigation canals into a single polylines layer. Roads and LULC were discrete variables, whereas human population density and density of linear features were continuous landscape variables used to generate the resistance surfaces. More details can be found in Thatte et al., (2020) and Tyagi et al., (2022). Previous studies have highlighted that the spatial scale of a landscape variable can affect the species–landscape relationship (Jackson & Fahrig, 2015). To address this issue, we analysed the landscape variables at seven different resolutions (0.25, 0.5, 1, 2.5, 5, 10, 25 km) for continuous variables (Hpop and Linden) and one categorical variable (LULC). While optimising the resistance landscape, a buffer of 20 km around all sample points was also incorporated.

#### 2.9.2. Multi-model optimization

To evaluate how landscape variables affect animal movement, we utilized a multi-model optimization approach to determine the maximum resistance offered (Rmax) and the relationship (x) between each variable and resistance. Genetic distance based on the proportion of shared alleles, D_PS_ (Bowcock et al., 1994), was used as the response variable. We used different parameter values for Rmax and x to identify the best combination that explained the genetic data. Seven values of maximum resistance (Rmax= 2, 5, 10, 50, 100, 500, 1000) and shape parameter (x = 0.01, 0.1, 0.5, 1, 2, 5, 10) were used. We calculated the cost distance for each pair of individuals for each combination of Rmax and x and utilized the MLPE.lmm() function from the *ResistanceGA* package (Peterman, 2018) to fit Linear mixed-effects (LME) models with maximum likelihood population effects (MLPE) parameterization. We identified the best model for each landscape variable based on the lowest AIC score from the MLPE mixed effects model (Shirk et al., 2010). In univariate optimization, we determined the scale and combination of parameters (Rmax and x) of the best-supported model for each landscape variable.

Best-fitting univariate models were combined additively and further optimized in a multivariate framework. We individually varied the model parameters (Rmax and x) for each landscape variable while holding the other variables’ parameter values constant (Thatte et al., 2020; Tyagi et al., 2022). If the optimal parameters changed for a landscape variable in the multivariate context, we held the new model parameters constant and varied the parameters of the next variable. We repeated this process until the parameterization of all the variables stabilized. This multivariate optimization approach enabled us to estimate resistance offered by each variable in combination with others for both species (Shirk et al., 2010).

### 2.10. Bootstrapping of landscape analysis

We used the resist.boot() function for bootstrapping within the *ResistanceGA* package (Peterman, 2018), with 10,000 iterations. Bootstrapping was performed using all four univariate optimization models, all the best multivariate combination models, and a null model for both species (Tables 3 and 4).

## 3. Results

### 3.1. Data summary

A total of 1,144 gaur and 756 sambar fecal samples (based on morphology) were collected from the field, and post-genetic species identity, 921 samples were identified as gaur, while 504 samples were determined to be of sambar. Subsequently, all identified samples underwent hDNA quantification before being selected for MBD enrichment. We selected 356 gaur and 288 sambar samples for MBD enrichment. After enrichment, the samples were quantified again, and those yielding a hDNA concentration of more than 0.06 ng/µl were selected for the preparation of ddRAD libraries (Tyagi et al., 2022). 171 gaur samples and 120 sambar samples were chosen for library preparation and sequencing, along with positive controls, extraction controls, PCR and library prep controls. The landscape and individual locations are shown in Figure 1.

### 3.2. Sequencing and SNP calling

Post demultiplexing and quality filtering, the average number of reads obtained per sample was 10.21 million for gaur (n=171) and 6.98 million for sambar (n=120). The higher number of reads in gaur samples was due to the data being sequenced twice and subsequently merged per sample (Fig S1). Mapping to the reference genome showed a higher percentage in sambar (average of 44.56%) compared to gaur (average of 32.63%) (Fig S1 and S2). Sample-wise detailed breakdown can be found in supplementary Tables S5 and S6.

The raw VCF file for gaur contained 30,47,450 variant sites, while for sambar, the number of variant sites in the raw VCF was 41,19,472. Post filtering, we excluded 47 gaur and 21 sambar samples, respectively due to high missing data. Individual identification was performed using relatedness values, and no recaptures were found in our dataset (Fig S3 and S4). The final putatively neutral dataset included 124 gaur individuals genotyped at 1186 SNPs and 2909 SNPs for 99 sambar individuals.

### 3.3. Genetic diversity, Isolation by distance and spatial autocorrelation

The genetic diversity of gaur was found to be low, with observed heterozygosity (Ho) of 0.129 and expected heterozygosity (He) of 0.149. In comparison, sambar also exhibited an overall low genetic diversity (Ho = 0.191; He = 0.22), although higher than that of gaur. Population-wise heterozygosity did not vary significantly for gaur, while there was variation observed for sambar (Table 1). Notably, both species exhibited a significant positive correlation between genetic and Euclidean distance, according to Mantel’s test (Gaur: Mantel’s R = 0.112, p-value = 0.004; Sambar: Mantel’s R = 0.117, p-value = 0.001; see Fig 2c and 3c). Spatial autocorrelation analysis revealed a significant positive spatial correlation up to 65 km and 30 km for gaur and sambar respectively (Fig 2d and 3d).

**Figure 2:**
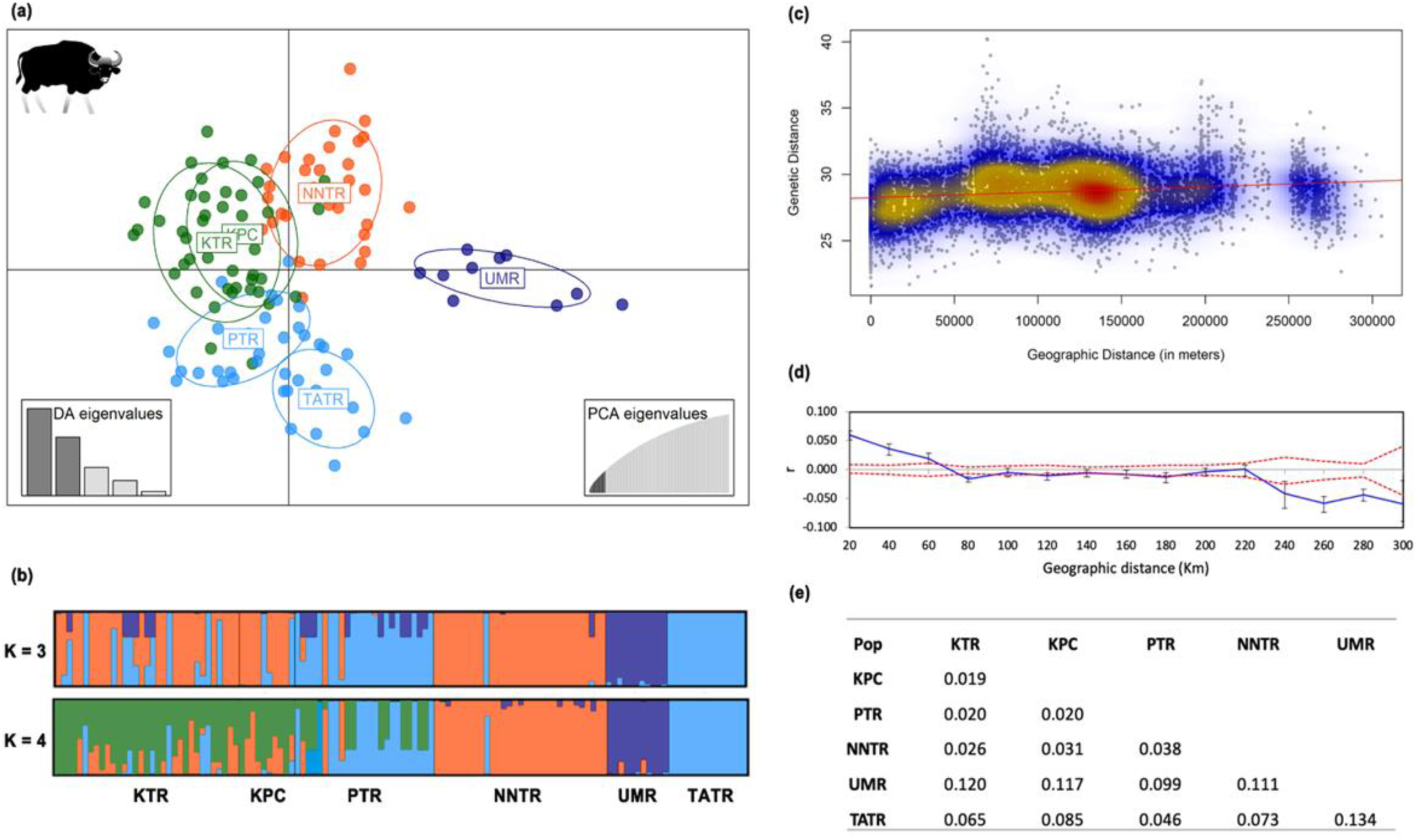
Population structure results of gaur described by (a) discriminant analysis of principal components (DAPC) based on 1186 SNPs. Each point represents a unique individual and each color corresponds to population, as assessed by STRUCTURE analysis. (b) STRUCTURE analysis plot where each vertical line represents a unique individual. Samples are colour-coded based on population. (c) Scatterplots for isolation by distance between genetic and geographic distance, colours indicate the density of data points as high (red), medium (yellow), and low (blue). (d) Spatial autocorrelation at distance classes of 20 km each till 300 km with r being the autocorrelation coefficient. Red dotted lines represent a 95% confidence interval. (e) Pairwise Fst between all the populations sampled.

**Figure 3:**
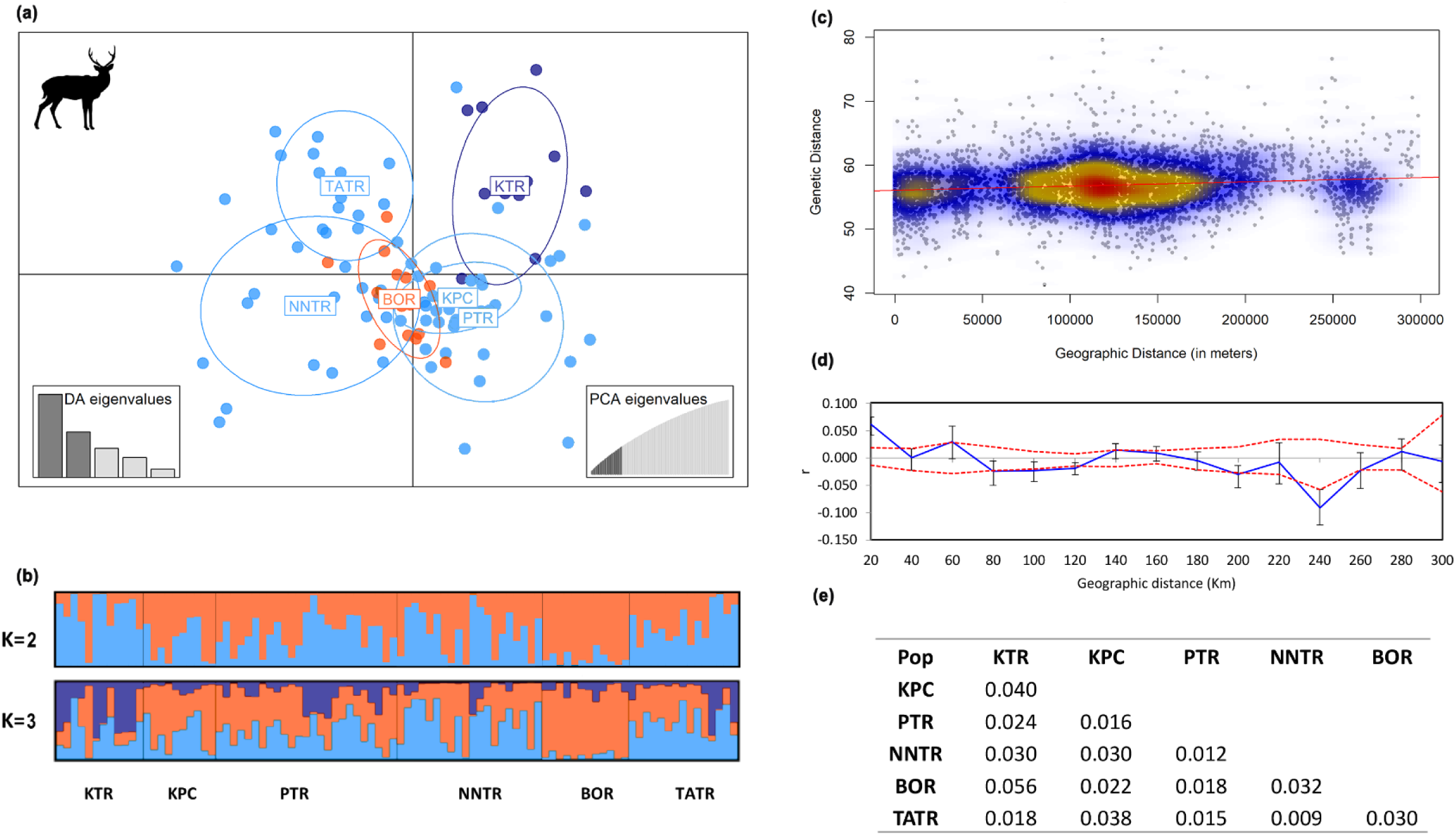
Population structure results of sambar based on (a) DAPC and (b) STRUCTURE analyses. Samples are colour-coded based on population. (c) Scatterplot between genetic and geographic distance. The density of data points is depicted with colours: red (high) to blue (low). (d) Spatial autocorrelation at distance classes of 20 km each till 300 km with r being autocorrelation coefficient. The dotted red lines represent a 95% confidence interval. (e) Pairwise Fst between all the populations sampled.

**Table 1:** Population-wise summary of the number of individuals, observed and expected heterozygosity for both species.

| Population | Gaur |  |  | Sambar |  |  |
| --- | --- | --- | --- | --- | --- | --- |
|  | N | Ho | He | N | Ho | He |
| KTR | 33 | 0.131 | 0.152 | 12 | 0.195 | 0.218 |
| KPC | 10 | 0.131 | 0.142 | 11 | 0.194 | 0.218 |
| PTR | 25 | 0.121 | 0.145 | 26 | 0.194 | 0.226 |
| NNTR | 31 | 0.124 | 0.152 | 21 | 0.187 | 0.226 |
| TATR | 14 | 0.129 | 0.147 | 15 | 0.209 | 0.224 |
| BOR | – | – | – | 14 | 0.165 | 0.209 |
| UMR | 11 | 0.122 | 0.148 | – | – | – |

### 3.4. Population genetic structure

Discriminant Analysis of Principal Components (DAPC) was conducted using the optimal number of PCs. For gaur, 14 PCs were retained based on the optim.a.score, revealing differentiation between Umred Karhandala Wildlife Sanctuary (UMR) and all other populations. Tadoba-Andhari Tiger Reserve (TATR) and Nagzira Nawegaon Tiger Reserve (NNTR) also exhibited differentiation (Fig 2a). STRUCTURE analyses indicated that the optimal number of genetic clusters for gaur was K = 3 (Fig 2b and S7). At K = 2, NNTR and UMR were segregated from the remaining populations, while at K = 3, UMR and NNTR formed separate clusters. At K = 4, UMR, NNTR, and TATR formed distinct clusters, while KTR, KPC, and PTR clustered together, displaying admixture (Fig 2b). Pairwise Fst also supported the results obtained from both DAPC and STRUCTURE analyses (Fig 2e). The highest pairwise Fst value was observed between UMR and TATR (0.134), whereas the lowest was found between KTR and KPC (0.019). UMR exhibited the highest pairwise Fst with all other populations, suggesting significant genetic differentiation and limited gene flow (Fig 2e).

In the case of sambar, a low level of genetic differentiation was observed. DAPC analysis using 22 PCs revealed overlap among all populations (Fig 3a). The most supported K value in STRUCTURE analysis was K = 2 (Evanno’s method, Fig 3b and S7). Pairwise Fst analysis also indicated low differentiation among sambar populations in central India (Fig 3e). The highest pairwise Fst value was observed between BOR and KTR (0.056), while the lowest was between NNTR and TATR (0.009).

### 3.5. Landscape connectivity and resistance analysis

Land use land cover (LULC) and roads with traffic were identified as the most influential factors impacting the genetic connectivity of both species. However, the resistance offered by different land use classes and traffic intensities were found to be different, as were the strengths of the correlation.

The impact of landscape modification on gaur movement was more pronounced than sambar, as indicated by the higher model fit (Table 2). In univariate analysis, all four landscape variables impact connectivity. Univariate analysis identified a spatial resolution of 25 km for Linden and Hpop, whereas LULC had the highest impact at a scale of 1 km (Table S2). LULC, roads with traffic, and the density of linear features exhibited the greatest resistance to gaur movement based on our multivariate analysis. Human population density did not contribute significantly to observed resistance, based on the best-fit model of multivariate analysis (Table 2; Fig 4). Moderate to high traffic roads offer high resistance to gaur movement. Areas with high linear feature density offered maximum resistance. Resistance increased as land cover changed to agriculture and human habitation (Fig S5). The selected model exhibited a marginal R^2^ of 0.503, explaining around 50% of the variation in the data (Table 2).

**Figure 4:**
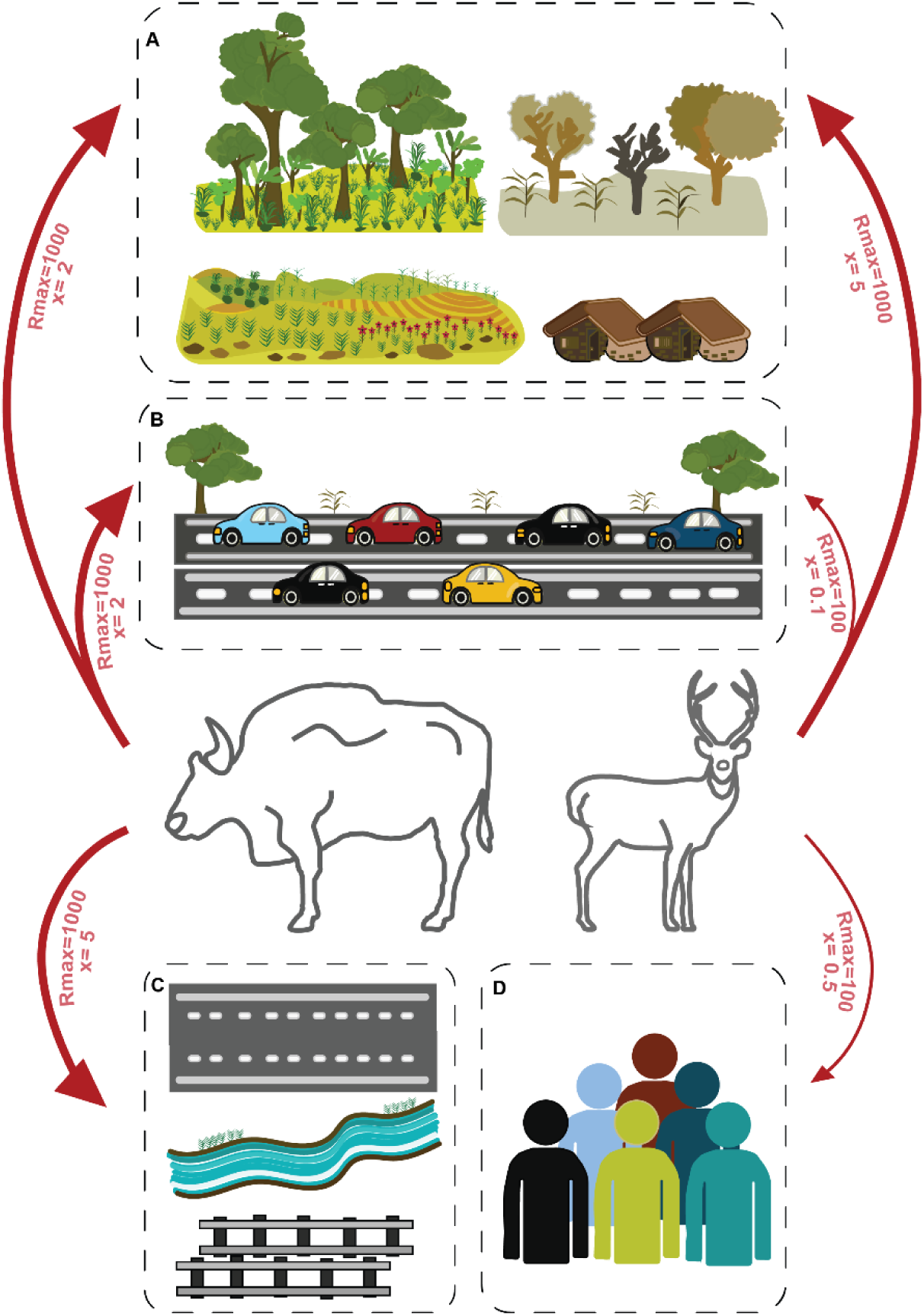
Association between landscape variables and resistance offered along with the optimized model parameters, Rmax and x, for both species. A- depicts land use and land cover variable, whereas B- depicts roads with traffic. The density of linear features and human population is depicted as C and D respectively. The thickness of the arrows shows the magnitude of resistance offered by each landscape variable.

**Table 2:** Multivariate optimization results for both gaur and sambar. The optimum parameters, Rmax: maximum resistance; x: shape parameter, for the different landscape variables for the best model based on lowest AIC score. The AIC score of the model in comparison with AIC score of the null model (Euclidian distance) along with the marginal R^2^ values are presented.

|  | Gaur |  |  |  |  | Sambar |  |  |  |  |
| --- | --- | --- | --- | --- | --- | --- | --- | --- | --- | --- |
| Layers | Rmax | x | AIC | AIC null | mR<sup>2</sup> | Rmax | x | AIC | AIC null | mR<sup>2</sup> |
| LULC | 1000 | 2 | -48431 | -47114.2 | 0.503 | 1000 | 5 | -24681 | -24627.2 | 0.173 |
| Roads | 1000 | 2 |  |  |  | 100 | 0.1 |  |  |  |
| Linden | 1000 | 5 |  |  |  | 50 | 5 |  |  |  |
| Hpop | 50 | 10 |  |  |  | 100 | 0.5 |  |  |  |

Univariate analysis for sambar (when landscape variables were tested individually) revealed that roads, human population density, and linear infrastructure density impacted connectivity, followed by land cover (Table S2). Specifically, roads strongly influenced sambar movement (x = 0.5; Table S2). Resistance demonstrated a linear increase with higher human population density (x = 1), and the shape parameter for linear feature density indicated that areas with high density offered greater resistance (Fig S6). Univariate analysis identified a spatial resolution of 25 km as optimal for all three layers (LULC, Hpop, Linden), suggesting their significance at a broader scale for sambar movement and connectivity. After combining the layers based on univariate optimization, the maximum resistance offered by the variables changed. In the final multivariate model, LULC emerged as the most influential variable for explaining connectivity, followed by roads with traffic and human population density (Table 2; Fig 4 & S5). However, the model fit was relatively low (mR^2^ = 0.173). Final resistance maps for both species were created using QGIS (Version 3.22.3) by employing optimized resistance values (Fig 5).

**Figure 5:**
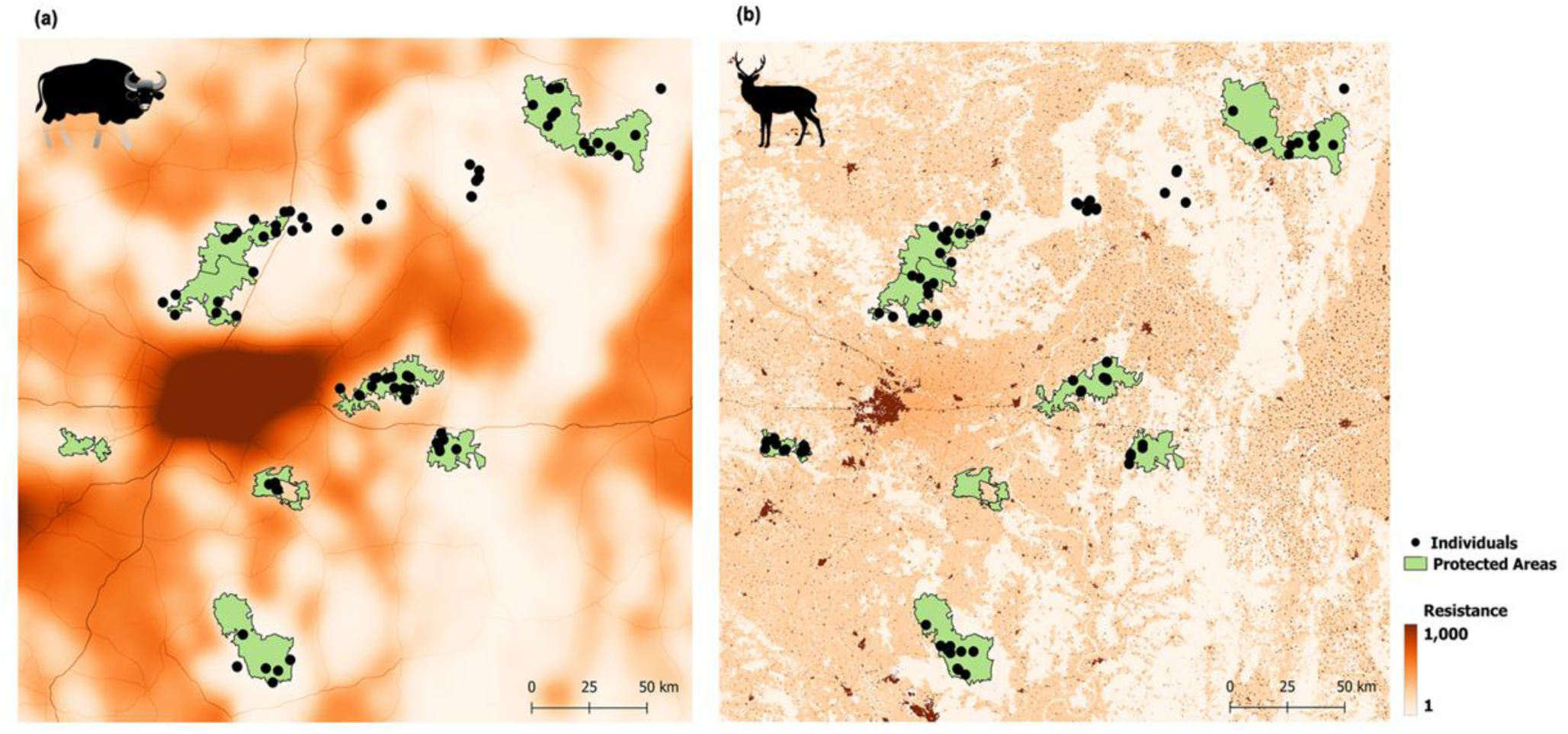
Optimized resistance surfaces for (a) gaur and (b) sambar. Dark colour represents areas with high resistance and light colours depict areas with low resistance. Black dots are the sample locations of the respective species on each map with protected areas marked as green polygons.

Bootstrapping analysis for gaur, with best models for all the possible combinations (n=15) along with the null model (Euclidian distance), indicated that the model consisting of Roads, LULC, and Linden achieved the highest mean log-likelihood score and had a selection probability of approximately 65% (Table 3). For sambar, the selection probability of the top model (including LULC, Roads, and Hpop) was 38%, followed by models incorporating Hpop and LULC, and one including Roads and Hpop (Table 4).

**Table 3:** Bootstrapping results for gaur with 15 models including all possible combinations of covariates along with the null (no resistance) model. The average AIC scores for the respective model is reported along with average weight, average rank and the average log- likelihood of the parameter combination. Percent top represents how many times the respective model was selected as the best model during 10,000 iterations.

| S.No. | Model with landscape variable combinations | Average AIC | Average weight | Average rank | Average LL | Percent top |
| --- | --- | --- | --- | --- | --- | --- |
| 1 | Roads+Linden+LULC | -27095.87 | 0.38 | 1.84 | 13550.93 | 65.76 |
| 2 | Hpop+Linden+LULC | -27095.76 | 0.36 | 2.39 | 13550.88 | 19.75 |
| 3 | Roads+Linden+LULC+Hpop | -27093.85 | 0.13 | 3.67 | 13550.93 | 0 |
| 4 | Roads+Linden | -27046.11 | 0.11 | 5.03 | 13525.06 | 11.86 |
| 5 | Linden | -27065.30 | 0.01 | 5.65 | 13533.65 | 0.96 |
| 6 | Hpop+LULC | -27029.47 | 0.01 | 7.17 | 13516.73 | 0.66 |
| 7 | Roads+Hpop+Linden | -27029.97 | 0.00 | 7.68 | 13517.99 | 0.01 |
| 8 | Roads+Linden | -27036.87 | 0.00 | 8.43 | 13520.44 | 0.06 |
| 9 | Roads+Hpop+LULC | -27027.46 | 0.00 | 8.49 | 13516.73 | 0 |
| 10 | Hpop+LULC | -27024.18 | 0.01 | 9.35 | 13514.09 | 0.9 |
| 11 | LULC | -27024.24 | 0.00 | 9.63 | 13513.12 | 0 |
| 12 | Roads+Hpop | -27021.94 | 0.00 | 10.22 | 13512.97 | 0.04 |
| 13 | Linden+LULC | -27003.70 | 0.00 | 12.22 | 13503.85 | 0 |
| 14 | Null | -26982.23 | 0.00 | 13.77 | 13491.12 | 0 |
| 15 | Roads | -26965.79 | 0.00 | 15.17 | 13483.89 | 0 |
| 16 | Hpop | -26965.22 | 0.00 | 15.27 | 13483.61 | 0 |

**Table 4:** Bootstrapping results for sambar with 15 models including all possible combinations of covariates along with the null (no resistance) model. The average AIC scores for the respective model is reported. The average weight, rank and the mean log- likelihood of the parameter combination are also shown along with how many times the respective model was selected as the best model during 10,000 iterations.

| S.No. | Model with landscape variable combinations | Average AIC | Average weight | Average rank | Average LL | Percent top |
| --- | --- | --- | --- | --- | --- | --- |
| 1 | Roads+Hpop+LULC | -13723.30 | 0.34 | 2.36 | 6863.65 | 38.6 |
| 2 | Hpop+LULC | -13723.10 | 0.30 | 2.51 | 6863.53 | 28 |
| 3 | Roads+Hpop | -13719.30 | 0.15 | 3.90 | 6863.66 | 18.1 |
| 4 | Roads+Hpop+LULC+Linden | -13718.80 | 0.05 | 4.79 | 6862.42 | 1.2 |
| 5 | LULC+Linden | -13708.50 | 0.05 | 6.60 | 6856.27 | 11.2 |
| 6 | Hpop+Linden | -13710.60 | 0.01 | 6.75 | 6857.29 | 0.8 |
| 7 | Roads+LULC | -13708.50 | 0.05 | 7.46 | 6856.27 | 0.4 |
| 8 | Hpop+Linden+LULC | -13711.10 | 0.01 | 7.68 | 6858.54 | 0 |
| 9 | Roads+Hpop+Linden | -13708.60 | 0.00 | 8.69 | 6857.29 | 0 |
| 10 | Roads+Linden+LULC | -13706.50 | 0.02 | 9.30 | 6856.27 | 0 |
| 11 | Roads | -13705.10 | 0.01 | 9.44 | 6853.55 | 0.7 |
| 12 | Roads+Hpop | -13705.10 | 0.00 | 9.70 | 6854.53 | 0.1 |
| 13 | LULC+Linden | -13698.30 | 0.01 | 11.84 | 6849.15 | 0.9 |
| 14 | Linden | -13670.00 | 0.00 | 14.50 | 6835.99 | 0 |
| 15 | Hpop | -13668.80 | 0.00 | 15.13 | 6835.41 | 0 |
| 16 | Null | -13668.70 | 0.00 | 15.36 | 6835.34 | 0 |

## 4. Discussion

Despite over two decades of research on connectivity, little is known about how large herbivores respond to various landscape features (Thatte et al., 2021). Addressing this gap, our study investigated the population structure and functional connectivity of gaur and sambar in central India. Using a customized ddRAD sequencing protocol for poor-quality DNA, we provide novel insights into genetic structure, impact of landscape features, and comparative connectivity of the two species.

Through a comprehensive analysis of thousands of genome-wide SNPs, we reveal distinct population structure for each species. Our landscape genetics analyses demonstrated that land use and linear infrastructure like roads hinder animal movement, highlighting differential species responses to landscape features. We identify small, isolated populations of both species in the landscape, and recommend that these need targeted conservation and management attention. Furthermore, our findings shed light on species-specific interactions with landscape elements, potentially influenced by life history traits and current population attributes, shaping spatial genetic structures. Such comparative datasets are important for understanding differential species response enabling holistic landscape-level conservation planning. Critically, these insights required the implementation of novel protocol optimizations.

### 4.1. ddRAD using fecal samples

We collected a substantial number of fecal samples for both species to mitigate loss due to poor DNA quality and quantity (Khan & Tyagi, 2021). Working with fecal DNA of herbivores can be challenging due to plant metabolites acting as PCR inhibitors, potentially limiting downstream processing for NGS data generation (Schrader et al., 2012). While we also observed sample loss at the enrichment stage, our ddRAD library preparation protocol minimized loss post-sequencing (due to insufficient data generation or missing data), an advancement over already available methods from previous studies (Tyagi et al., 2022).

Our protocol is robust and cost-effective. With the given protocol and designed indexes, 468 samples can be pooled and sequenced on any Illumina platform (in our case HiSeq2500, but can also be sequenced on NextSeq or NovaSeq). Most of the available ddRAD protocols are designed to use around 200 ng of DNA as starting material (Magbanua et al., 2023; Peterson et al., 2012; Peterson et al., 2014). We used as little as 5 ng of enriched host DNA for data generation. Next-generation sequencing has allowed creative approaches to access genome-wide or genotyping data from a wide array of non-invasive samples (including fecal samples). Such approaches include SNP panels, capture baits, and enrichment methods, among others (Chiou & Bergey, 2018; Hayward et al., 2022; Khan et al., 2020; Natesh et al., 2019; Orkin et al., 2020; Snyder-Mackler et al., 2016; Tyagi et al., 2022). Our study adds to the growing body of literature enhancing conservation genomics application for wildlife studies and significantly improving applicability, cost and DNA concentration requirements.

### 4.2. Population genetic structure

Our study spotlights low genetic diversity in gaur and sambar populations, underscoring habitat degradation and fragmentation threats. We document low diversity and high population differentiation among gaur populations in central India, which aligns with previous findings from another protected area in the same region using mitochondrial and microsatellite data (Atkulwar et al., 2020; Farah et al., 2021). Our results highlight that the small population of Umred-Karhandala Wildlife Sanctuary (UMR) is the most genetically differentiated, despite being geographically central to the three large protected areas (Fig 1 and Fig 2a & b). This pattern could be driven by isolation and genetic drift, as this is the smallest population among the sampled populations. In STRUCTURE analysis, at higher K values of 3 and 4, populations like NNTR and TATR also separate into distinct clusters (Fig 2b). This was unlike KTR, KPC and PTR, where we always observed admixture. This could result from a well-maintained corridor (KPC) between these two protected areas (KTR and PTR). Many studies have highlighted the importance and functionality of this corridor using data from different species like tiger (Schoen et al., 2022; Thatte et al., 2018), leopard (Dutta et al., 2013) and sloth bear (Dutta et al., 2015). Our study highlights the significance and functionality of existing corridors in maintaining connectivity for large herbivores like gaur.

However, sambar populations exhibit low differentiation and genetic diversity (relatively higher as compared to gaur). This contrasts with our expectations based on the species’ similar habitat requirements. A previous study on sambar in the western Himalayas, also reported similar genetic diversity using mitochondrial and microsatellite datasets (Singh et al., 2021). The population in Bor Tiger Reserve, which is close to cities and separated by national highways (Fig 1), displays the highest Fst value and the least genetic diversity (Fig 3b & 3e; Table 1), highlighting limited animal movement. Overall, low genetic differentiation in sambar might suggest that this species can utilize a variety of land covers to disperse across the landscape.

Both species showed a weak but significant correlation between genetic distance and geographical distance (Fig 2c and 3c). The spatial autocorrelation relationship broke down at distances beyond ∼65 km for gaur and ∼30 km for sambar, which is in agreement with dispersal abilities of these species. The weak correlation indicated that isolation-by-distance has a weak effect, and other factors might be more important in shaping the population structure. For example, the geographic distance between KTR and TATR is around 300 km, while that between UMR and TATR is around 100 km, but pairwise Fst is higher for the later pair of populations (0.065 versus 0.134 respectively). This highlights dispersal limitation caused by landscape factors other than geographic distance, also documented in various other species (Rengefors et al., 2021; Thatte et al., 2020).

### 4.3. Impact of anthropogenic disturbance on functional connectivity

Land use land cover (LULC) and roads with traffic emerged as key factors influencing genetic connectivity for both species. While agriculture and human habitation greatly impacted gaur movement, agriculture had a variable impact on sambar gene flow. This aligns with connectivity trends observed in other large mammals like tiger, leopard (Thatte et al., 2020) and mule deer (Fraser et al., 2019). High density of linear features (roads, railway tracks, canals) had high resistance for gaur movement, known to influence gene flow of other species like roe deer (Coulon et al., 2006), and can potentially lead to genetic differentiation (Thatte et al., 2020; Weckworth et al., 2013).

Numerous studies have shown that road networks affect the connectivity and hinder gene flow of various wildlife populations (Epps et al., 2005; Ernest et al., 2014; Fraser et al., 2019; Laurance & Balmford, 2013; Thatte et al., 2018, 2020; Tyagi et al., 2022). Our findings reveal varying impacts of road networks and traffic intensities on our study species. Medium to high traffic roads significantly affect gaur movement in the landscape, with resistance increasing non-linearly (x=2), and was highest with high traffic. In contrast, sambar experiences lower resistance on small and low-traffic roads compared to gaur. Roads seem to impact large herbivores (gaur and sambar) more severely than large carnivores like tigers (Thatte et al., 2020), which are primarily affected by high-traffic roads rather than low to mid-traffic intensities. India has the second largest road network globally and central India has a complex road network with varying traffic intensities. Our results indicate that while these networks may not act as absolute barriers to movement for species like tigers, they have a significant negative impact on herbivores like gaur, among others (suggested in Thatte et al., 2020). This emphasizes the need for wildlife connectivity mitigation measures on smaller roads in addition to major state and national highways.

Human population density was found to have a low impact on gaur movement in our analysis (maybe due to forest-restricted habitat association) but impacts sambar movement significantly in the landscape. The magnitude of resistance offered was not as high as LULC, but was similar to the road networks (Rmax = 100). Our findings using genome-wide data resonate with a previous study in the same landscape (Jayadevan et al., 2020), which suggested that human land use, population density, and linear infrastructure could hinder both species’ movement. Our study provides a more detailed analysis and empirical evidence supporting these landscape responses.

The univariate analyses indicated differing optimal scales for the same landscape variable between the two species. The optimal LULC scale for gaur was 1 km, while for sambar, it was 25 km. In contrast, both species were impacted by human population density and linear feature density at a 25 km scale. This distinct response highlights gaur’s susceptibility to minor habitat modifications and landscape changes, whereas sambar appears more resilient to small-scale alterations in land use patterns.

Our multivariate optimization approach offered comprehensive insight into the combined impacts of multiple landscape variables on genetic connectivity. Landscape genetic analysis is particularly reliable for species or populations with significant genetic differentiation and when landscape variables strongly affect gene flow (Shirk et al., 2017, 2018). In central India, gaur populations exhibit high differentiation, with our analysis demonstrating a significant role of landscape in shaping genetic structure. The model fit was strong, with an mR^2^ of 0.503. Bootstrapping analysis supports the robustness of our findings, confirming the best-fit model’s selection probability at approximately 65%.

However, for sambar, the genetic differentiation is low, possibly due to a large effective population size and wide distribution, mitigating detectable genetic structure (Whitlock & McCauley, 1999). Potential past translocations may also contribute to reduced genetic structure. Our landscape genetics results imply lower landscape impact on sambar movement. The model fit for sambar was lower (mR^2^ = 0.173), with bootstrapping findings suggesting selection probabilities for multiple models instead of a single best-fit model (Table 2 & 4; Fig 4). After 10,000 iterations, the top model had a selection probability of around 38% (Table 4). Lower model fit may not always imply lack of impact of landscape on connectivity, but also the inability of the methods to detect those. More samples from the broader geographic range or an even higher number of SNPs might improve landscape genetic inferences for sambar. This underscores the need to develop novel methods for assessing landscape feature impact on species with low-genetic-differentiation. There are a few other caveats worth mentioning here, including the need for good quality reference genomes for effective conservation of endangered species (Paez et al., 2022). In our study, we utilized the reference genomes of closely related species due to the unavailability of high-quality reference assembly for both of our study species. Another crucial point for future studies adopting our approach is the quality and quantity of field samples. We recommend collecting fresh, and a higher number of samples to mitigate potential losses during enrichment, library preparation, and filtering.

### 4.4. Conservation implications

This study presents the first landscape-wide connectivity assessment for large herbivores using genomic tools in India. Our methods and models can be applied to other data-deficient endangered species in other landscapes. Urban areas and road networks are expected to grow substantially by 2030 (Seto et al., 2012), and human populations are also expected to increase, especially in biodiversity hotspots (Cincotta et al., 2000). The combined effects of human demographic growth, land use change, and climate change pose a serious threat to wildlife connectivity globally (Sanderson et al., 2019). In India, infrastructure development like the expansion of cities, construction of new roads and expansion of existing highways (http://morth.nic.in/BharatMala), railway networks and interlinking river projects, and construction of canals and dams, are prominent threats to wild species and their habitats. Despite the need for development to meet India’s economic goals, it is imperative to align development with conservation objectives. While the Wildlife Protection Act of India restricts major developments in and around Tiger Reserves (which are designated as eco-sensitive zones), the provisions for construction and implementation of mitigation measures on roads and railway lines, which bisect crucial wildlife movement areas, are limited, partly due to lack of data on animal movement. Our results tried to bridge this gap by providing information on how different species with similar habitat associations can respond to various landscape alterations.

Central India is a priority conservation landscape and stronghold of multiple endangered species (Jhala et al., 2011), but is exposed to anthropogenic pressures like land use change, mining, and road network expansion. Along with this wildlife also faces pressures like human-wildlife conflict, tourism pressure, risk of hybridization and disease transfer from domestic species (Miller et al., 2015; Srivathsa et al., 2019; Tyagi et al., 2019, 2023). While most connectivity focused on single charismatic species like tigers (Schoen et al., 2022; Thatte et al., 2018; Yumnam et al., 2014), we highlight that it is crucial to consider the connectivity requirements of different species in the region, as they exhibit varied responses to landscape features based on their habitat associations and requirements. This could eventually aid in the conservation and maintenance of populations for top carnivores such as tigers. Carnivore populations might not solely rely on their connectivity but may also depend on connected, viable populations of wild prey species. In the future, detailed understanding of priority species coupled with broader scale attributes (e.g., threats and habitats) will help with on-ground landscape prioritization and species management, and aid in conservation planning.

In conclusion, our study contributes to the growing body of literature on landscape genetics and the conservation of large herbivores in the face of anthropogenic pressures using cutting-edge genomic and landscape genetic methods. We identified common and different responses of two study species to landscape elements. Our work identified priority areas, including smaller populations like UMR and BOR, vital for gene flow and restoration, which could help secure the long-term survival of these species. Our work lays the groundwork for holistic conservation and prioritization strategies by combining genetic data with landscape resistance. Our results offer valuable insights for wildlife managers and policymakers, to mitigate the impacts of habitat loss, fragmentation, and impending development on large mammals, to enhance biodiversity management.

## Author contributions

Conceptualization: A.T., and U.R., Data collection: A.T., Laboratory work: A.T., N.Y., and A.P., Data analyses: A.T., Project administration: A.T., N.Y., and U.R., Data Curation: A.T., N.Y., and A.P., Supervision: A.T. and U.R., Funding acquisition: A.T. and U.R., Writing - Original Draft: A.T., Writing - Review & Editing: A.T., N.Y., A.P., and U.R.

## Acknowledgements

We thank the forest departments of Maharashtra (No: Desk-22(8)/CR-12(17-18)/928/2017-2018, dated 29/6/2017 and Desk-22(8)/WL/research/ CR-12(17 -18)/2660/20-21/2^nd^ February 2021) and Madhya Pradesh (S.No./Technical-I/5104, dated: 27/07/2017 and S.No./M.CH-II/Research/7130, dated: 26/11/2020) for their support in granting the necessary permits and logistics during the sample collection. We would also like to thank the dedicated forest department staff and our field assistants for their steady assistance throughout the sample collection phase. We extend our thanks to all the remarkable volunteers, namely Varun Taneja, Basawaraj Mulage, Tanvi Gurjar, Himani Khangwal, Sourav Mehra, Hrishikesh Wagh, Varad Bansod, Shardul Joshi, Purnima Mohite, Abhilasha Fulzele, Abhishek Shukla, Srishti Manna, Neha More, Ashish Thoke, Mrunali Rout, Akshay Narnaware, and Saurav Jagtap for their immense efforts and support during the fieldwork, which was instrumental for the sample collection. We thank Ankit Kacchwaha from The Corbett Foundation for his invaluable help in organizing logistics and providing support throughout the fieldwork. We also thank Tejali Naik and Gokul Nair for their invaluable contributions during the data generation. We thank Abishek Harihar and Divya Vasudev for providing crucial comments on the manuscript. We would like to acknowledge the Wildlife Conservation Trust (WCT) for generously providing a small grant to A.T., which not only supported the fieldwork but also aided in preliminary laboratory analysis, including DNA extractions. Additionally, we acknowledge The Defries Bajpai Foundation (DBF) for awarding a small grant to A.T., which covered the second field season. For institutional support, we thank National Centre for Biological Sciences (NCBS). Furthermore, we would like to acknowledge the Next Generation Genomics Facility (NGGF, Bangalore Life Science Cluster, BLiSC) for assistance during data generation. NCBS data cluster used is supported under project no. 12-R&D-TFR-5.04-0900, Department of Atomic Energy, Government of India. AT was supported by NCBS and the Council for Industrial and Scientific Research-University Grant Commission fellowship.

## Data availability statement

Raw demultiplexed sequence data for both the species, generated in this study has been made available on NCBI Sequence Read Archive (SRA) database and can be found under the BioProject accession numbers PRJNA1121501 and PRJNA1121523 for gaur and sambar respectively.

