## Supplementary information for "On the road to losing connectivity: Fecal samples provide genome-wide insights into anthropogenic impacts on two large herbivore species in central India"

MOLECULAR ECOLOGY

**Table of Contents:**

| S.No. | Description | Page number |
|---|---|---|
| 1 | **S1:** Species-specific primer designing and PCR conditions used | 3 |
| 2 | **Table S1:** Details of species-specific primers designed in this study | 4 |
| 3 | **Table S2:** Univariate optimization results for both gaur and sambar | 4 |
| 4 | **Table S3:** Details of adapters and modified sequencing primers designed for ddRAD sequencing in this study | 5 |
| 5 | **Table S4:** Details of indexes used for ddRAD in this study | 5-6 |
| 6 | **Table S5:** Sample-wise details for all gaur samples | 7-11 |
| 7 | **Table S6:** Sample-wise details for all sambar samples | 12-14 |
| 8 | **Figure S1:** Reads obtained and mapped | 15 |
| 9 | **Figure S2:** Mapping percentage | 16 |
| 10 | **Figure S3:** Relatedness values for all gaur individuals | 17 |
| 11 | **Figure S4:** Relatedness values for all sambar individuals | 18 |
| 12 | **Figure S5:** Rmax and x for the best model for Gaur | 19 |
| 13 | **Figure S6:** Rmax and x for the best model for Sambar | 20 |
| 14 | **Figure S7:** Plots for estimating the optimal value of K | 21 |
| 15 | **S2:** ddRAD protocol designed in this study | 22-29 |
| 16 | **References** | 30 |

**S1: Species-specific primer designing and PCR conditions used**.

To design species-specific primers for Gaur and Sambar, we aligned cytochrome b gene of mitochondrial genome of both these species with other sympatric species with which the fecal samples could be misidentified based on morphology. For designing gaur specific primer, we used mithun (*Bos frontalis*), gaur (*Bos gaurus*), zebu cattle (*Bos indicus*), domestic cattle (*Bos taurus*), buffalo (Bubalus bubalis), wild water buffalo (Bubalus arnee), nilgai (*Boselaphus tragocamelus*), Asiatic elephant (*Cervus elaphus*), Indian rhinoceros (*Rhinoceros unicornis*), chital (*Axis axis*), swamp deer (*Rucervus duvaucelii*), sambar (*Rusa unicolor*), Indian muntjac (*Muntiacus muntjak*), blackbuck (*Antilope cervicapra*), Indian gazelle (*Gazella bennettii*), hangul (*Cervus hanglu*), brow-antlered deer (*Rucervus eldii*), hog deer (*Axis porcinus*), four-horned antelope (*Tetracerus quadricornis*), sloth bear (*Melursus ursinus*), wild boar (*Sus scrofa*) and humans (*Homo sapiens*).

To develop primers that are specific to Sambar, we aligned cytochrome b gene of four-horned antelope (*Tetracerus quadricornis*), Indian muntjac (*Muntiacus muntjak*), domestic goat (*Capra hircus*), nilgiri tahr (*Nilgiritragus hylocrius*), nilgai (Boselaphus tragocamelus), chital (Axis axis), sambar (*Rusa unicolor*), swamp deer (Rucervus duvaucelii), hog deer (Axis porcinus), Indian gazelle (*Gazella bennettii*), blackbuck (*Antilope cervicapra*), rhesus macaque (*Macaca mulatta*), grey langur (*Semnopithecus entellus*), wild boar (*Sus scrofa*) and humans (*Homo sapiens*). Primer sequences can be found in Table S1. Our gaur-specific primer was unable to distinguish between gaur and Mithun (*Bos frontalis*), due to very high similarity in cytochrome b gene for both species.

PCR amplifications were performed in 10 µL reaction volumes, consisting of 5 µL of 2x Multiplex PCR master mix (Qiagen), 0.5 µL of each primer (2 µM), and 2 µL of DNA template and 2 µM of PCR grade water. The PCR cycling conditions involved an initial denaturation step at 95 °C for 15 min, followed by 40 cycles of denaturation at 95 °C for 30 sec, annealing at 52 °C for 30 sec, and extension at 72 °C for 30 sec. The final extension step was set at 72 °C for 10 mins.

**Table S1:** Details of species-specific primers designed in this study. The primer names, sequence, amplicon size and annealing temperatures are provided.

| Species | Primer name | Sequence 5'-3' | Amplicon (bp) | Tm (°C) |
|---|---|---|---|---|
| Gaur | Bg_cytb1.1F | CACAGCAATTGCCATAGTCC | 151 | 52 |
| | Bg_cytb1.1R | AAGGGCTAGAATTAGTAACAAGGTT | | |
| Sambar | CommF_CytB1 | ATTGTAAACAACGCATTCAT | 115 | 52 |
| | SambarR_CytB1 | GGAATAGGCCTGTGATG | | |

**Table S2:** Univariate optimization results for both gaur and sambar. The table reports optimum parameters (Rmax—maximum resistance offered and x—shape parameter) for all the landscape variables. The AIC value of the best-fit model along with AIC value of the null model is reported. The optimum spatial resolution for land use land cover (LULC), density of linear features and human population density is also reported.

| | Gaur | | | | | Sambar | | | | |
|---|---|---|---|---|---|---|---|---|---|---|
| **Layers** | **Rmax** | **x** | **AIC** | **AIC null** | **Scale (m)** | **Rmax** | **x** | **AIC** | **AIC null** | **Scale (m)** |
| **LULC** | 1000 | 0.5 | -47241.3 | -47114.2 | 1000 | 50 | 5 | -24681 | -24627.2 | 25000 |
| **Roads** | 1000 | 0.5 | -47137.9 | | NA | 1000 | 0.5 | -24642.7 | | NA |
| **Linden** | 1000 | 5 | -47270.1 | | 25000 | 1000 | 10 | -24671.1 | | 25000 |
| **Hpop** | 1000 | 1 | -47131.5 | | 25000 | 1000 | 1 | -24637.5 | | 25000 |

**Table S3:** Details of adapters and modified sequencing primers designed for ddRAD library preparation and sequencing on HiSeq2500 run in this study.

| Name | Oligo Sequence (5'-3') |
|---|---|
| **SphI_Adapter 1A** | ACACTCTTTCCCTACACGACGCTCTTCCGATCTCATG |
| **SphI_Adapter 1B** | /5Phos/AGATCGGAAGAGCGTCGTGTAGGGAAAGAGTGT |
| **MluCI_Adapter 2A** | GTGACTGGAGTTCAGACGTGTGCTCTTCCGATCT |
| **MluCI_Adapter 2B** | /5Phos/AATTAGATCGGAAGAGCGAGAACAA |
| **Modified R1_SPH1** | ACACTCTTTCCCTACACGACGCTCTTCCGATCTCATGC |
| **Modified R2_Mluc1** | GTGACTGGAGTTCAGACGTGTGCTCTTCCGATCTAATT |

**Table S4:** Details of indexes used for ddRAD library preparation in this study. Index sequences are adapted from Illumina *Nextera* Index sequences (468 combinations). Unique barcode sequences are highlighted.

| Name | Oligo Sequence (5'-3') | Bases in Adapter |
|---|---|---|
| **i7-01** | CAAGCAGAAGACGGCATACGAGATTCGCCTTAGTGACTGGAGTTCAGACGTGTGC | TCGCCTTA |
| **i7-02** | CAAGCAGAAGACGGCATACGAGATCTAGTACGGTGACTGGAGTTCAGACGTGTGC | CTAGTACG |
| **i7-03** | CAAGCAGAAGACGGCATACGAGATTTCTGCCTGTGACTGGAGTTCAGACGTGTGC | TTCTGCCT |
| **i7-04** | CAAGCAGAAGACGGCATACGAGATGCTCAGGAGTGACTGGAGTTCAGACGTGTGC | GCTCAGGA |
| **i7-05** | CAAGCAGAAGACGGCATACGAGATAGGAGTCCGTGACTGGAGTTCAGACGTGTGC | AGGAGTCC |
| **i7-06** | CAAGCAGAAGACGGCATACGAGATCATGCCTAGTGACTGGAGTTCAGACGTGTGC | CATGCCTA |
| **i7-07** | CAAGCAGAAGACGGCATACGAGATGTAGAGAGGTGACTGGAGTTCAGACGTGTGC | GTAGAGAG |
| **i7-08** | CAAGCAGAAGACGGCATACGAGATCCTCTCTGGTGACTGGAGTTCAGACGTGTGC | CCTCTCTG |

| | | |
|---|---|---|
| **i7-09** | CAAGCAGAAGACGGCATACGAGATAGCGTAGCGTGACTGGAGTTCAGACGTGTGC | AGCGTAGC |
| **i7-10** | CAAGCAGAAGACGGCATACGAGATCAGCCTCGGTGACTGGAGTTCAGACGTGTGC | CAGCCTCG |
| **i7-11** | CAAGCAGAAGACGGCATACGAGATTGCCTCTTGTGACTGGAGTTCAGACGTGTGC | TGCCTCTT |
| **i7-12** | CAAGCAGAAGACGGCATACGAGATTCCTCTACGTGACTGGAGTTCAGACGTGTGC | TCCTCTAC |
| **i7-14** | CAAGCAGAAGACGGCATACGAGATTCATGAGCGTGACTGGAGTTCAGACGTGTGC | TCATGAGC |
| **i7-15** | CAAGCAGAAGACGGCATACGAGATCCTGAGATGTGACTGGAGTTCAGACGTGTGC | CCTGAGAT |
| **i7-16** | CAAGCAGAAGACGGCATACGAGATTAGCGAGTGTGACTGGAGTTCAGACGTGTGC | TAGCGAGT |
| **i7-18** | CAAGCAGAAGACGGCATACGAGATGTAGCTCCGTGACTGGAGTTCAGACGTGTGC | GTAGCTCC |
| **i7-19** | CAAGCAGAAGACGGCATACGAGATTACTACGCGTGACTGGAGTTCAGACGTGTGC | TACTACGC |
| **i7-20** | CAAGCAGAAGACGGCATACGAGATAGGCTCCGGTGACTGGAGTTCAGACGTGTGC | AGGCTCCG |
| **i7-21** | CAAGCAGAAGACGGCATACGAGATGCAGCGTAGTGACTGGAGTTCAGACGTGTGC | GCAGCGTA |
| **i7-22** | CAAGCAGAAGACGGCATACGAGATCTGCGCATGTGACTGGAGTTCAGACGTGTGC | CTGCGCAT |
| **i7-23** | CAAGCAGAAGACGGCATACGAGATGAGCGCTAGTGACTGGAGTTCAGACGTGTGC | GAGCGCTA |
| **i7-24** | CAAGCAGAAGACGGCATACGAGATCGCTCAGTGTGACTGGAGTTCAGACGTGTGC | CGCTCAGT |
| **i7-26** | CAAGCAGAAGACGGCATACGAGATGTCTTAGGGTGACTGGAGTTCAGACGTGTGC | GTCTTAGG |
| **i7-27** | CAAGCAGAAGACGGCATACGAGATACTGATCGGTGACTGGAGTTCAGACGTGTGC | ACTGATCG |
| **i7-28** | CAAGCAGAAGACGGCATACGAGATTAGCTGCAGTGACTGGAGTTCAGACGTGTGC | TAGCTGCA |
| **i7-29** | CAAGCAGAAGACGGCATACGAGATGACGTCGAGTGACTGGAGTTCAGACGTGTGC | GACGTCGA |
| **i5-01** | AATGATACGGCGACCACCGAGATCTACACTAGATCGCACACTCTTTCCCTACACGACG | TAGATCGC |
| **i5-02** | AATGATACGGCGACCACCGAGATCTACACCTCTCTATACACTCTTTCCCTACACGACG | CTCTCTAT |
| **i5-03** | AATGATACGGCGACCACCGAGATCTACACTATCCTCTACACTCTTTCCCTACACGACG | TATCCTCT |
| **i5-04** | AATGATACGGCGACCACCGAGATCTACACAGAGTAGAACACTCTTTCCCTACACGACG | AGAGTAGA |
| **i5-05** | AATGATACGGCGACCACCGAGATCTACACGTAAGGAGACACTCTTTCCCTACACGACG | GTAAGGAG |
| **i5-06** | AATGATACGGCGACCACCGAGATCTACACACTGCATAACACTCTTTCCCTACACGACG | ACTGCATA |
| **i5-07** | AATGATACGGCGACCACCGAGATCTACACAAGGAGTAACACTCTTTCCCTACACGACG | AAGGAGTA |
| **i5-08** | AATGATACGGCGACCACCGAGATCTACACCTAAGCCTACACTCTTTCCCTACACGACG | CTAAGCCT |
| **i5-10** | AATGATACGGCGACCACCGAGATCTACACCGTCTAATACACTCTTTCCCTACACGACG | CGTCTAAT |
| **i5-11** | AATGATACGGCGACCACCGAGATCTACACTCTCTCCGACACTCTTTCCCTACACGACG | TCTCTCCG |
| **i5-13** | AATGATACGGCGACCACCGAGATCTACACTCGACTAGACACTCTTTCCCTACACGACG | TCGACTAG |
| **i5-15** | AATGATACGGCGACCACCGAGATCTACACTTCTAGCTACACTCTTTCCCTACACGACG | TTCTAGCT |
| **i5-16** | AATGATACGGCGACCACCGAGATCTACACCCTAGAGTACACTCTTTCCCTACACGACG | CCTAGAGT |
| **i5-17** | AATGATACGGCGACCACCGAGATCTACACGCGTAAGAACACTCTTTCCCTACACGACG | GCGTAAGA |
| **i5-18** | AATGATACGGCGACCACCGAGATCTACACCTATTAAGACACTCTTTCCCTACACGACG | CTATTAAG |
| **i5-20** | AATGATACGGCGACCACCGAGATCTACACAAGGCTATACACTCTTTCCCTACACGACG | AAGGCTAT |
| **i5-21** | AATGATACGGCGACCACCGAGATCTACACGAGCCTTAACACTCTTTCCCTACACGACG | GAGCCTTA |
| **i5-22** | AATGATACGGCGACCACCGAGATCTACACTTATGCGAACACTCTTTCCCTACACGACG | TTATGCGA |

**Table S5:** Sample-wise details for gaur: details of reads obtained, mapped and mapping percentage for all the sequenced gaur samples with populations are included (124 samples included in the analysis). **Samples excluded from the analysis post filtering the data.

| S.NO. | SAMPLE | POP | COMBINED RUN1 + RUN2 | | |
|---|---|---|---|---|---|
| | | | READS OBTAINED | READS MAPPED | MAPPING %AGE |
| **1** | NNTR897 | NNTR | 9467412 | 2816134 | 29.75 |
| **2** | NNTR931 | NNTR | 9855514 | 2581794 | 26.20 |
| **3** | TATR584 | TATR | 9340995 | 2824306 | 30.24 |
| **4** | TATR599 | TATR | 15486619 | 5231517 | 33.78 |
| **5** | TATR600 | TATR | 11414757 | 3689027 | 32.32 |
| **6** | TATR602 | TATR | 10172324 | 3234514 | 31.80 |
| **7** | TATR603 | TATR | 9981188 | 2591377 | 25.96 |
| **8** | TATR619 | TATR | 12337829 | 3450754 | 27.97 |
| **9** | TATR630 | TATR | 12316698 | 3696510 | 30.01 |
| **10** | TATR697 | TATR | 11818075 | 3784371 | 32.02 |
| **11** | TATR698 | TATR | 9679749 | 2783893 | 28.76 |
| **12** | TATR699 | TATR | 8902414 | 2367990 | 26.60 |
| **13** | TATR700 | TATR | 8174000 | 2404070 | 29.41 |
| **14** | TATR704 | TATR | 7184338 | 1815424 | 25.27 |
| **15** | TATR800 | TATR | 9279565 | 2450981 | 26.41 |
| **16** | TATR801 | TATR | 8448610 | 2534968 | 30.00 |
| **17** | WCT1084 | PTR | 12705060 | 4869392 | 38.33 |
| **18** | WCT1091 | PTR | 13340001 | 4370329 | 32.76 |
| **19** | WCT1098 | PTR | 10783595 | 3773967 | 35.00 |
| **20** | WCT124 | KTR | 9793251 | 2765195 | 28.24 |
| **21** | WCT1265 | PTR | 11930763 | 4026175 | 33.75 |
| **22** | WCT1272 | PTR | 4886266 | 1632341 | 33.41 |
| **23** | WCT1296 | PTR | 7706621 | 2371247 | 30.77 |
| **24** | WCT1300 | PTR | 17351707 | 6432149 | 37.07 |
| **25** | WCT1347 | NNTR | 12815591 | 4547925 | 35.49 |
| **26** | WCT1348 | NNTR | 14022880 | 4792732 | 34.18 |
| **27** | WCT1349 | NNTR | 15584795 | 5535048 | 35.52 |
| **28** | WCT1355 | NNTR | 12731597 | 4805446 | 37.74 |
| **29** | WCT1359 | NNTR | 6994115 | 2683892 | 38.37 |
| **30** | WCT1363 | NNTR | 3533362 | 1210331 | 34.25 |
| **31** | WCT1409 | NNTR | 14155701 | 4491258 | 31.73 |
| **32** | WCT1436 | NNTR | 13928848 | 4092505 | 29.38 |
| **33** | WCT1445 | NNTR | 8599185 | 2493791 | 29.00 |
| **34** | WCT1447 | NNTR | 11738076 | 3686629 | 31.41 |

| | | | | | |
|---|---|---|---|---|---|
| **35** | WCT1448 | NNTR | 10084403 | 3349555 | 33.22 |
| **36** | WCT1455 | NNTR | 14793274 | 5050338 | 34.14 |
| **37** | WCT1457 | NNTR | 11153591 | 3521212 | 31.57 |
| **38** | WCT1498 | NNTR | 8523358 | 2447238 | 28.71 |
| **39** | WCT1509 | NNTR | 11923153 | 3724522 | 31.24 |
| **40** | WCT1510 | NNTR | 11083356 | 2921297 | 26.36 |
| **41** | WCT1527 | NNTR | 8844257 | 2824480 | 31.94 |
| **42** | WCT1579 | NNTR | 13420565 | 5432290 | 40.48 |
| **43** | WCT1590 | NNTR | 13334419 | 5599643 | 41.99 |
| **44** | WCT1591 | NNTR | 11184961 | 3887911 | 34.76 |
| **45** | WCT1627 | NNTR | 7526302 | 2697327 | 35.84 |
| **46** | WCT1631 | NNTR | 12276109 | 4097132 | 33.37 |
| **47** | WCT1671 | NNTR | 5207312 | 1610469 | 30.93 |
| **48** | WCT1673 | NNTR | 11912449 | 5173533 | 43.43 |
| **49** | WCT1680 | NNTR | 17454296 | 7701793 | 44.13 |
| **50** | WCT1710 | NNTR | 14097893 | 5399918 | 38.30 |
| **51** | WCT1725 | NNTR | 15235403 | 5302772 | 34.81 |
| **52** | WCT1747 | NNTR | 12634861 | 4460874 | 35.31 |
| **53** | WCT1759 | NNTR | 12861338 | 4606067 | 35.81 |
| **54** | WCT1791 | UMR | 13934703 | 5783452 | 41.50 |
| **55** | WCT1797 | UMR | 15035116 | 8985300 | 59.76 |
| **56** | WCT1800 | UMR | 13807298 | 4770401 | 34.55 |
| **57** | WCT1801 | UMR | 11472599 | 3945073 | 34.39 |
| **58** | WCT1802 | UMR | 13422433 | 4562352 | 33.99 |
| **59** | WCT1803 | UMR | 14652284 | 6337417 | 43.25 |
| **60** | WCT1804 | UMR | 15327524 | 4439002 | 28.96 |
| **61** | WCT1805 | UMR | 13949647 | 5936447 | 42.56 |
| **62** | WCT1807 | UMR | 15313754 | 6022118 | 39.32 |
| **63** | WCT1811 | UMR | 12754952 | 6432933 | 50.43 |
| **64** | WCT1817 | UMR | 15902463 | 6988174 | 43.94 |
| **65** | WCT188 | KTR | 4107275 | 1490683 | 36.29 |
| **66** | WCT201 | KTR | 11915136 | 3772213 | 31.66 |
| **67** | WCT203 | KTR | 9762856 | 2545439 | 26.07 |
| **68** | WCT21 | KTR | 10788557 | 4585630 | 42.50 |
| **69** | WCT217 | KTR | 11330712 | 3920685 | 34.60 |
| **70** | WCT219 | KTR | 12872337 | 4045009 | 31.42 |
| **71** | WCT225 | KTR | 13512894 | 4533405 | 33.55 |
| **72** | WCT258 | KTR | 12297580 | 4359965 | 35.45 |
| **73** | WCT260 | KTR | 10572554 | 3190776 | 30.18 |
| **74** | WCT262 | KTR | 10260302 | 3235102 | 31.53 |
| **75** | WCT267 | KTR | 11738748 | 3708476 | 31.59 |
| **76** | WCT269 | KTR | 11220816 | 3553349 | 31.67 |
| **77** | WCT273 | KTR | 10968205 | 3396595 | 30.97 |

| | | | | | |
|---|---|---|---|---|---|
| **78** | WCT277 | KTR | 13199398 | 4512419 | 34.19 |
| **79** | WCT279 | KTR | 16041638 | 5694535 | 35.50 |
| **80** | WCT294 | KTR | 15988203 | 5556939 | 34.76 |
| **81** | WCT295 | KTR | 13440771 | 4214402 | 31.36 |
| **82** | WCT297 | KTR | 13567311 | 4360169 | 32.14 |
| **83** | WCT300 | KTR | 12386100 | 3872117 | 31.26 |
| **84** | WCT301 | KTR | 11130751 | 3356434 | 30.15 |
| **85** | WCT302 | KTR | 10849697 | 3153647 | 29.07 |
| **86** | WCT304 | KTR | 10802068 | 3466229 | 32.09 |
| **87** | WCT329 | KTR | 12328077 | 3559536 | 28.87 |
| **88** | WCT330 | KTR | 11502507 | 3508890 | 30.51 |
| **89** | WCT332 | KTR | 13180524 | 4272224 | 32.41 |
| **90** | WCT333 | KTR | 10829343 | 3502683 | 32.34 |
| **91** | WCT335 | KTR | 14885860 | 4980555 | 33.46 |
| **92** | WCT339 | KTR | 14629034 | 4679951 | 31.99 |
| **93** | WCT355 | KTR | 13113306 | 3378703 | 25.77 |
| **94** | WCT382 | KTR | 5442803 | 1704808 | 31.32 |
| **95** | WCT387 | KTR | 9702232 | 2942750 | 30.33 |
| **96** | WCT388 | KTR | 5284852 | 1512702 | 28.62 |
| **97** | WCT542 | KPC | 10280309 | 3527215 | 34.31 |
| **98** | WCT545 | KPC | 14629563 | 5300470 | 36.23 |
| **99** | WCT565 | KPC | 12002380 | 3370241 | 28.08 |
| **100** | WCT568 | KPC | 10916449 | 3968837 | 36.36 |
| **101** | WCT570 | KPC | 13258497 | 4650657 | 35.08 |
| **102** | WCT590 | KPC | 11656689 | 3216775 | 27.60 |
| **103** | WCT619 | KPC | 12190216 | 3985388 | 32.69 |
| **104** | WCT638 | KPC | 14535662 | 3827865 | 26.33 |
| **105** | WCT640 | KPC | 13115944 | 3134355 | 23.90 |
| **106** | WCT643 | KPC | 10859198 | 2584061 | 23.80 |
| **107** | WCT659 | PTR | 14371771 | 4774984 | 33.22 |
| **108** | WCT669 | PTR | 6441214 | 1995509 | 30.98 |
| **109** | WCT686 | PTR | 7338131 | 2389416 | 32.56 |
| **110** | WCT743 | PTR | 11429252 | 3674075 | 32.15 |
| **111** | WCT744 | PTR | 2993890 | 1025112 | 34.24 |
| **112** | WCT745 | PTR | 2921307 | 1058887 | 36.25 |
| **113** | WCT771 | PTR | 14245323 | 4635603 | 32.54 |
| **114** | WCT786 | PTR | 12010782 | 3632202 | 30.24 |
| **115** | WCT787 | PTR | 14888387 | 5549955 | 37.28 |
| **116** | WCT788 | PTR | 3181294 | 926743 | 29.13 |
| **117** | WCT804 | PTR | 3003151 | 1025451 | 34.15 |
| **118** | WCT811 | PTR | 7103959 | 2445615 | 34.43 |
| **119** | WCT815 | PTR | 9181243 | 2972195 | 32.37 |
| **120** | WCT840 | PTR | 6017789 | 1734076 | 28.82 |

| | | | | | |
|---|---|---|---|---|---|
| **121** | WCT862 | PTR | 12992525 | 5528672 | 42.55 |
| **122** | WCT917 | PTR | 11333453 | 3318233 | 29.28 |
| **123** | WCT951 | PTR | 10375405 | 2876867 | 27.73 |
| **124** | WCT965 | PTR | 10264903 | 3471993 | 33.82 |
| **125** | TATR707 | ** | 5440726 | 1370065 | 25.18 |
| **126** | TATR709 | ** | 6751487 | 1780843 | 26.38 |
| **127** | TATR712 | ** | 7053984 | 2144783 | 30.41 |
| **128** | TATR780 | ** | 9227294 | 2592894 | 28.10 |
| **129** | TATR783 | ** | 5242472 | 1274247 | 24.31 |
| **130** | TATR803 | ** | 9735237 | 2228819 | 22.89 |
| **131** | TATR805 | ** | 9212640 | 2691918 | 29.22 |
| **132** | WCT1001 | ** | 8720854 | 5851814 | 67.10 |
| **133** | WCT1003 | ** | 8457606 | 2480892 | 29.33 |
| **134** | WCT1008 | ** | 7164660 | 1970706 | 27.51 |
| **135** | WCT1086 | ** | 7673396 | 2591024 | 33.77 |
| **136** | WCT1128 | ** | 7423106 | 2133346 | 28.74 |
| **137** | WCT1158 | ** | 10025000 | 3078198 | 30.71 |
| **138** | WCT1160 | ** | 5950780 | 1435094 | 24.12 |
| **139** | WCT1194 | ** | 8049344 | 3010894 | 37.41 |
| **140** | WCT1231 | ** | 7320776 | 1612010 | 22.02 |
| **141** | WCT1243 | ** | 11905700 | 4407890 | 37.02 |
| **142** | WCT1245 | ** | 7241488 | 2253394 | 31.12 |
| **143** | WCT1247 | ** | 3214412 | 930366 | 28.94 |
| **144** | WCT1271 | ** | 3761280 | 1258840 | 33.47 |
| **145** | WCT1280 | ** | 7522626 | 2536732 | 33.72 |
| **146** | WCT1410 | ** | 9077802 | 2814712 | 31.01 |
| **147** | WCT1449 | ** | 6537364 | 1913334 | 29.27 |
| **148** | WCT1464 | ** | 8254170 | 2540432 | 30.78 |
| **149** | WCT1465 | ** | 4608486 | 1454810 | 31.57 |
| **150** | WCT147 | ** | 8078986 | 2325286 | 28.78 |
| **151** | WCT1484 | ** | 6756402 | 2252818 | 33.34 |
| **152** | WCT1491 | ** | 7345502 | 2635762 | 35.88 |
| **153** | WCT1503 | ** | 8334518 | 2746174 | 32.95 |
| **154** | WCT1504 | ** | 6757842 | 2084070 | 30.84 |
| **155** | WCT1782 | ** | 7097170 | 2535190 | 35.72 |
| **156** | WCT235 | ** | 5590342 | 1764312 | 31.56 |
| **157** | WCT239 | ** | 5378338 | 1667396 | 31.00 |
| **158** | WCT359 | ** | 7095892 | 2159546 | 30.43 |
| **159** | WCT380 | ** | 4539680 | 1281708 | 28.23 |
| **160** | WCT414 | ** | 6908244 | 2360560 | 34.17 |
| **161** | WCT420 | ** | 8121876 | 3051900 | 37.58 |
| **162** | WCT469 | ** | 8353294 | 2154250 | 25.79 |
| **163** | WCT660 | ** | 9324916 | 3378048 | 36.23 |

| | | | | | |
|---|---|---|---|---|---|
| **164** | WCT661 | ** | 9601114 | 3113958 | 32.43 |
| **165** | WCT663 | ** | 9327528 | 3219278 | 34.51 |
| **166** | WCT87 | ** | 6493060 | 1927146 | 29.68 |
| **167** | WCT871 | ** | 9083646 | 2300156 | 25.32 |
| **168** | WCT922 | ** | 6656978 | 1781160 | 26.76 |
| **169** | WCT938 | ** | 6492674 | 1906758 | 29.37 |
| **170** | WCT959 | ** | 7701484 | 2533518 | 32.90 |
| **171** | WCT981 | ** | 6957876 | 2115016 | 30.40 |

*Note: During filtering, we removed 209 loci for gaur based on Bayescan analysis.*

**Table S6:** Sample-wise details of sambar: details of reads obtained, mapped and mapping percentage for all the sequenced gaur samples with populations are included (99 samples included in the analysis). **Samples excluded from the analysis post filtering the data.

| S.NO. | SAMPLE | POP | READS OBTAINED | READS MAPPED | MAPPING %AGE |
|---|---|---|---|---|---|
| **1** | DBF477 | BOR | 6926865 | 2384018 | 34.42% |
| **2** | DBF482 | BOR | 9628704 | 3674846 | 38.17% |
| **3** | DBF484 | BOR | 10498899 | 6111468 | 58.21% |
| **4** | DBF487 | BOR | 8689712 | 2957628 | 34.04% |
| **5** | DBF490 | BOR | 7378622 | 2730387 | 37.00% |
| **6** | DBF499 | BOR | 3883873 | 1548417 | 39.87% |
| **7** | DBF500 | BOR | 8016294 | 2940213 | 36.68% |
| **8** | DBF502 | BOR | 7295001 | 2599804 | 35.64% |
| **9** | DBF510 | BOR | 6247066 | 2835582 | 45.39% |
| **10** | DBF516 | BOR | 7578194 | 2754742 | 36.35% |
| **11** | DBF518 | BOR | 6608996 | 2373097 | 35.91% |
| **12** | DBF527 | BOR | 6549302 | 2537815 | 38.75% |
| **13** | DBF530 | BOR | 8490580 | 3195313 | 37.63% |
| **14** | DBF531 | BOR | 7533984 | 2965029 | 39.36% |
| **15** | DBF826 | NNTR | 16224054 | 14266676 | 87.94% |
| **16** | DBF847 | NNTR | 12086594 | 6077243 | 50.28% |
| **17** | DBF850 | NNTR | 5512906 | 2974547 | 53.96% |
| **18** | DBF856 | NNTR | 8247750 | 4044799 | 49.04% |
| **19** | DBF867 | NNTR | 7871237 | 3658148 | 46.47% |
| **20** | DBF868 | NNTR | 7530465 | 3419582 | 45.41% |
| **21** | DBF884 | NNTR | 9756836 | 4492621 | 46.05% |
| **22** | DBF885 | NNTR | 8469041 | 4103887 | 48.46% |
| **23** | DBF886 | NNTR | 8699665 | 3511577 | 40.36% |
| **24** | DBF888 | NNTR | 8432614 | 3323343 | 39.41% |
| **25** | DBF894 | NNTR | 8810727 | 3520776 | 39.96% |
| **26** | DBF904 | NNTR | 10383592 | 4410624 | 42.48% |
| **27** | DBF906 | NNTR | 7553152 | 2892746 | 38.30% |
| **28** | DBF918 | NNTR | 10954701 | 9937663 | 90.72% |
| **29** | DBF949 | NNTR | 10970920 | 4305634 | 39.25% |
| **30** | DBF950 | NNTR | 8530996 | 5024266 | 58.89% |
| **31** | DBF952 | NNTR | 8990125 | 3989546 | 44.38% |
| **32** | DBF271 | PTR | 7532220 | 5636960 | 74.84% |
| **33** | DBF287 | PTR | 5374403 | 1842876 | 34.29% |
| **34** | DBF348 | PTR | 8616304 | 4281584 | 49.69% |
| **35** | DBF349 | PTR | 7642140 | 2819969 | 36.90% |

| | | | | | |
|---|---|---|---|---|---|
| **36** | DBF356 | PTR | 7658713 | 4046771 | 52.84% |
| **37** | DBF386 | PTR | 9746794 | 4225456 | 43.35% |
| **38** | DBF470 | PTR | 6008431 | 2108640 | 35.09% |
| **39** | DBF105 | PTR | 8522141 | 3133246 | 36.77% |
| **40** | DBF114 | PTR | 10683623 | 4942528 | 46.26% |
| **41** | DBF127 | PTR | 7082298 | 2425963 | 34.25% |
| **42** | DBF138 | PTR | 10108964 | 3568422 | 35.30% |
| **43** | DBF139 | PTR | 10092855 | 3930054 | 38.94% |
| **44** | DBF153 | PTR | 9763573 | 3645887 | 37.34% |
| **45** | DBF166 | PTR | 7485122 | 3932663 | 52.54% |
| **46** | DBF168 | PTR | 6816604 | 2259067 | 33.14% |
| **47** | DBF196 | PTR | 5768312 | 2226690 | 38.60% |
| **48** | DBF34 | PTR | 8242492 | 3478480 | 42.20% |
| **49** | DBF640 | TATR | 6680679 | 3061184 | 45.82% |
| **50** | DBF662 | TATR | 8582951 | 3199460 | 37.28% |
| **51** | DBF664 | TATR | 6377460 | 1720115 | 26.97% |
| **52** | DBF665 | TATR | 10171531 | 4455034 | 43.80% |
| **53** | DBF713 | TATR | 6595466 | 2843780 | 43.12% |
| **54** | DBF718 | TATR | 7169341 | 3516844 | 49.05% |
| **55** | DBF721 | TATR | 8598227 | 3755130 | 43.67% |
| **56** | DBF723 | TATR | 9376496 | 4557849 | 48.61% |
| **57** | DBF738 | TATR | 12872098 | 12829158 | 99.67% |
| **58** | DBF739 | TATR | 7668154 | 3049496 | 39.77% |
| **59** | DBF741 | TATR | 8509007 | 3936759 | 46.27% |
| **60** | DBF742 | TATR | 8841486 | 7565023 | 85.56% |
| **61** | DBF744 | TATR | 7575423 | 2916083 | 38.49% |
| **62** | DBF745 | TATR | 10032340 | 8499282 | 84.72% |
| **63** | DBF777 | TATR | 10421904 | 4690440 | 45.01% |
| **64** | WCT184 | KTR | 5651820 | 3068623 | 54.29% |
| **65** | WCT009 | KTR | 2329511 | 1189669 | 51.07% |
| **66** | WCT034 | KTR | 6892006 | 5746279 | 83.38% |
| **67** | WCT076 | KTR | 5442643 | 2499528 | 45.92% |
| **68** | WCT1052 | PTR | 6087191 | 2552968 | 41.94% |
| **69** | WCT1066 | PTR | 9175702 | 3156557 | 34.40% |
| **70** | WCT1193 | PTR | 10820610 | 5228210 | 48.32% |
| **71** | WCT1200 | PTR | 6696656 | 2489474 | 37.17% |
| **72** | WCT1318 | PTR | 8576839 | 3341403 | 38.96% |
| **73** | WCT1336 | PTR | 6775819 | 3202878 | 47.27% |
| **74** | WCT1439 | NNTR | 12408025 | 4666903 | 37.61% |
| **75** | WCT1442 | NNTR | 10025921 | 4482199 | 44.71% |
| **76** | WCT1534 | NNTR | 7099389 | 2566192 | 36.15% |
| **77** | WCT1692 | NNTR | 7307597 | 2712273 | 37.12% |
| **78** | WCT185 | KTR | 6007236 | 5783749 | 96.28% |

| | | | | | |
|---|---|---|---|---|---|
| **79** | WCT298 | KTR | 7050265 | 2656012 | 37.67% |
| **80** | WCT310 | KTR | 6187859 | 2437766 | 39.40% |
| **81** | WCT316 | KTR | 5018870 | 2257641 | 44.98% |
| **82** | WCT321 | KTR | 5022643 | 1666193 | 33.17% |
| **83** | WCT340 | KTR | 7479574 | 4961783 | 66.34% |
| **84** | WCT404 | KTR | 5998944 | 3189354 | 53.17% |
| **85** | WCT405 | KTR | 4690209 | 2087715 | 44.51% |
| **86** | WCT556 | KPC | 6442992 | 2402061 | 37.28% |
| **87** | WCT559 | KPC | 2552607 | 1072194 | 42.00% |
| **88** | WCT575 | KPC | 1850160 | 926217 | 50.06% |
| **89** | WCT577 | KPC | 5251577 | 2004866 | 38.18% |
| **90** | WCT591 | KPC | 1581277 | 801102 | 50.66% |
| **91** | WCT600 | KPC | 5478811 | 2658050 | 48.52% |
| **92** | WCT604 | KPC | 4901340 | 1694730 | 34.58% |
| **93** | WCT605 | KPC | 10463787 | 4024863 | 38.46% |
| **94** | WCT607 | KPC | 7753160 | 3108523 | 40.09% |
| **95** | WCT608 | KPC | 5932957 | 1758412 | 29.64% |
| **96** | WCT610 | KPC | 6633646 | 3074607 | 46.35% |
| **97** | WCT762 | PTR | 7214342 | 6933851 | 96.11% |
| **98** | WCT776 | PTR | 8385103 | 3496182 | 41.70% |
| **99** | WCT941 | PTR | 4914925 | 2485900 | 50.58% |
| **100** | DBF486 | ** | 8260323 | 2875905 | 34.82% |
| **101** | DBF496 | ** | 1705956 | 575447 | 33.73% |
| **102** | DBF106 | ** | 9768059 | 3267529 | 33.45% |
| **103** | DBF129 | ** | 9996368 | 3282895 | 32.84% |
| **104** | DBF32 | ** | 10262319 | 3681200 | 35.87% |
| **105** | DBF595 | ** | 2722542 | 951217 | 34.94% |
| **106** | WCT1354 | ** | 7314565 | 2904253 | 39.71% |
| **107** | WCT287 | ** | 7552969 | 2312641 | 30.62% |
| **108** | WCT299 | ** | 1816671 | 682772 | 37.58% |
| **109** | WCT320 | ** | 6824673 | 1631630 | 23.91% |
| **110** | WCT343 | ** | 5781134 | 2169769 | 37.53% |
| **111** | WCT354 | ** | 3719100 | 1152586 | 30.99% |
| **112** | WCT476 | ** | 6437797 | 3905079 | 60.66% |
| **113** | WCT558 | ** | 3504209 | 1272241 | 36.31% |
| **114** | WCT594 | ** | 2745788 | 1148029 | 41.81% |
| **115** | WCT595 | ** | 4550217 | 1430418 | 31.44% |
| **116** | WCT599 | ** | 7116361 | 2365379 | 33.24% |
| **117** | WCT614 | ** | 2516309 | 837365 | 33.28% |
| **118** | WCT615 | ** | 5294515 | 1844200 | 34.83% |
| **119** | WCT727 | ** | 6621577 | 2557540 | 38.62% |
| **120** | WCT761 | ** | 8168469 | 1920604 | 23.51% |

*Note: During filtering, we removed 631 loci for sambar based on Bayescan analysis.*

**Figure S1: Reads obtained and mapped.** Plot showing number of reads obtained and mapped to reference genome for both Gaur and Sambar. For Gaur, the combined number of reads was counted for both sequencing runs.

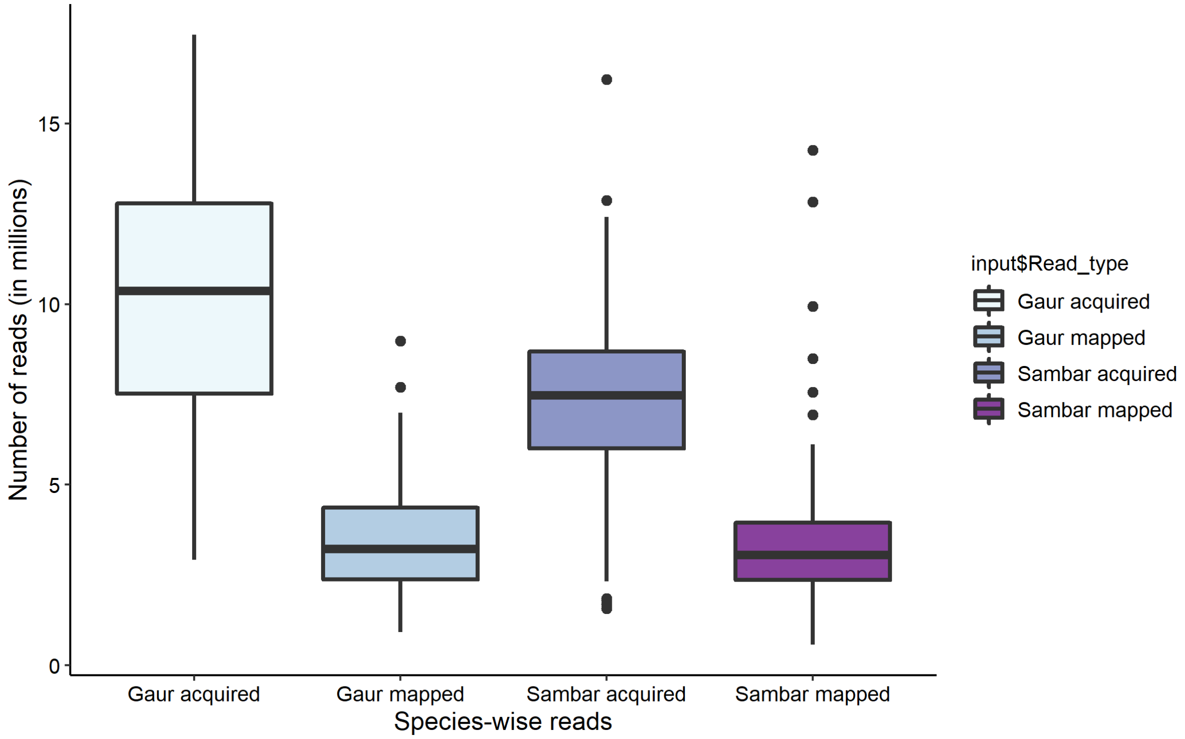

**Figure S2: Mapping percentage.** Plot showing mapping percentage of the obtained reads to the respective reference genomes. Gaur reads were mapped to cow reference genome (ARS-UCD1.2; RefSeq accession: GCF_002263795.1), whereas sambar reads were mapped to red deer reference assembly (mCerEla1.1; RefSeq accession: GCF_910594005.1).

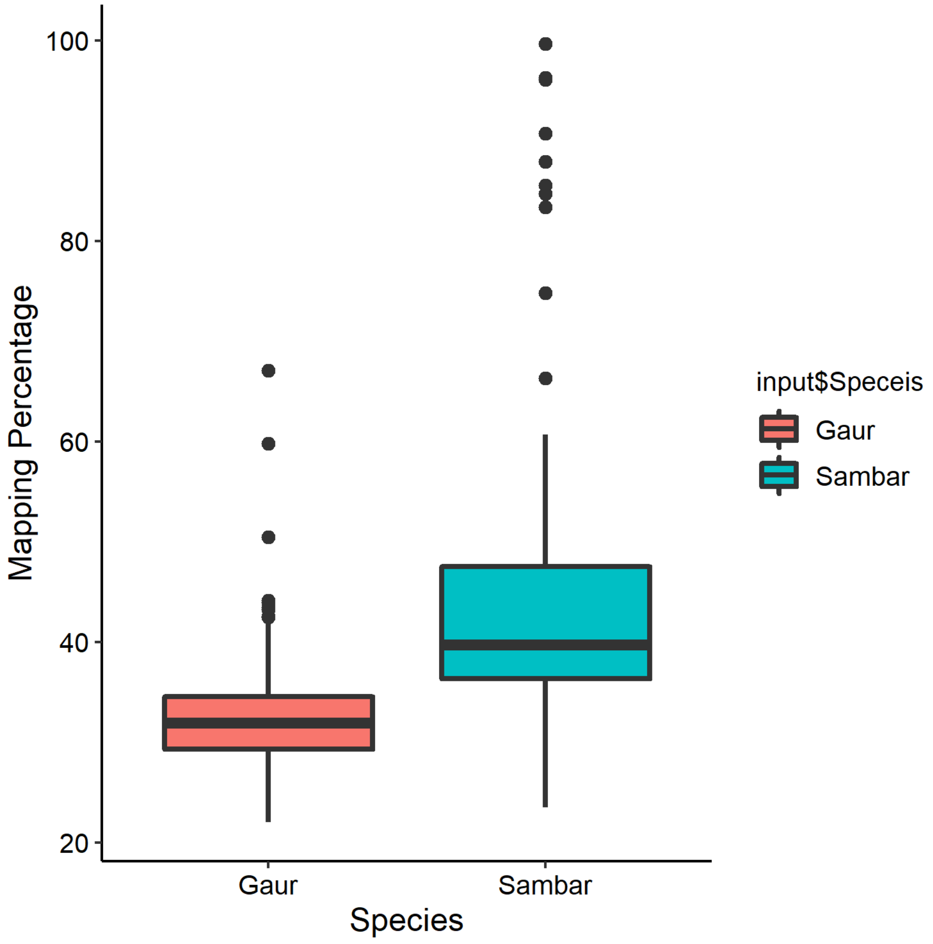

**Figure S3: Relatedness values for all gaur individuals.** Heat map of relatedness values obtained for all 124 gaur individuals with three relatedness bins (0-0.2; 0.2-0.4 and 0.8-1).

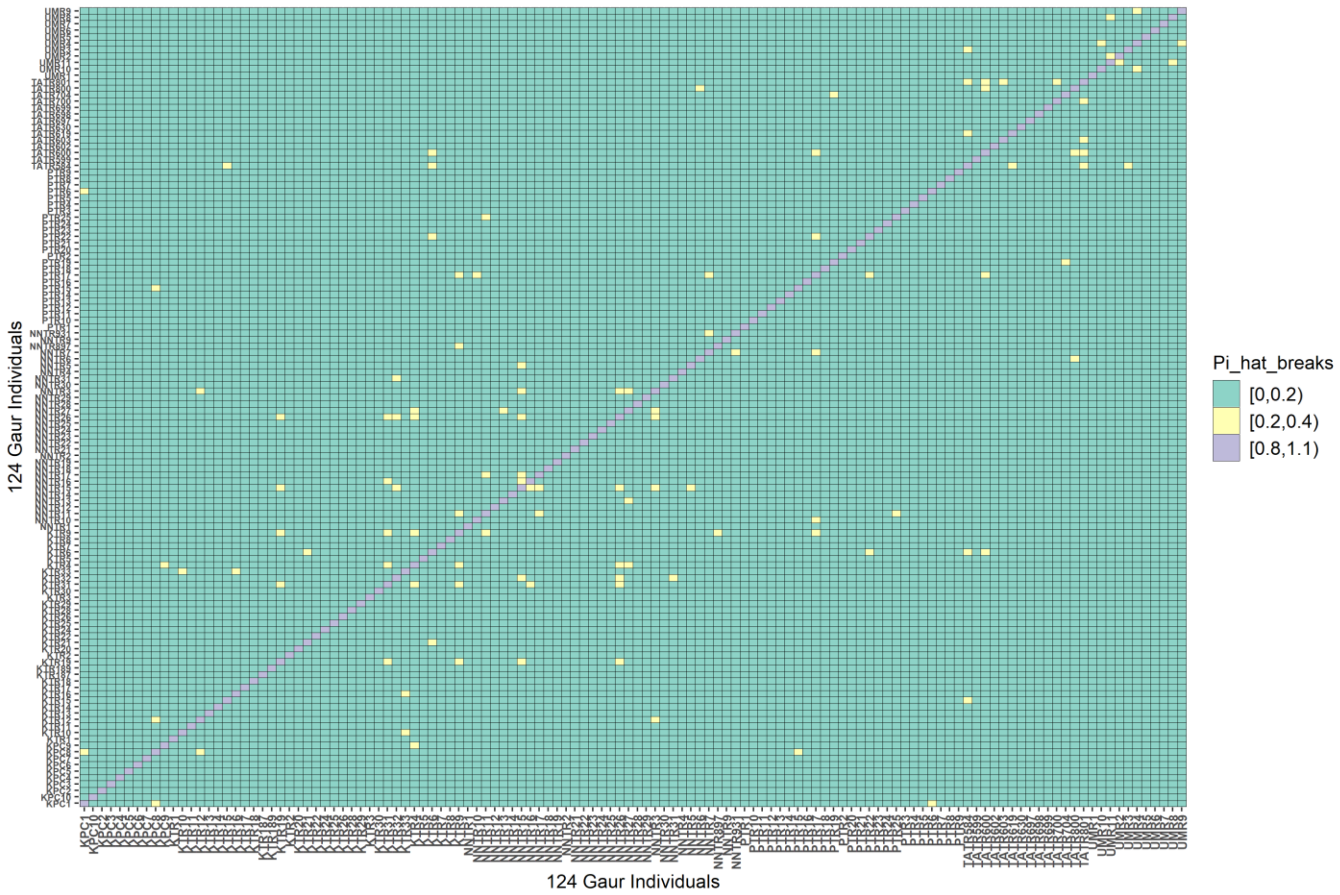

**Figure S4: Relatedness values for all sambar individuals.** Heat map of relatedness values obtained for all 99 Sambar individuals with four relatedness bins (0-0.2; 0.2-0.4, 0.4-0.6 and 0.8-1).

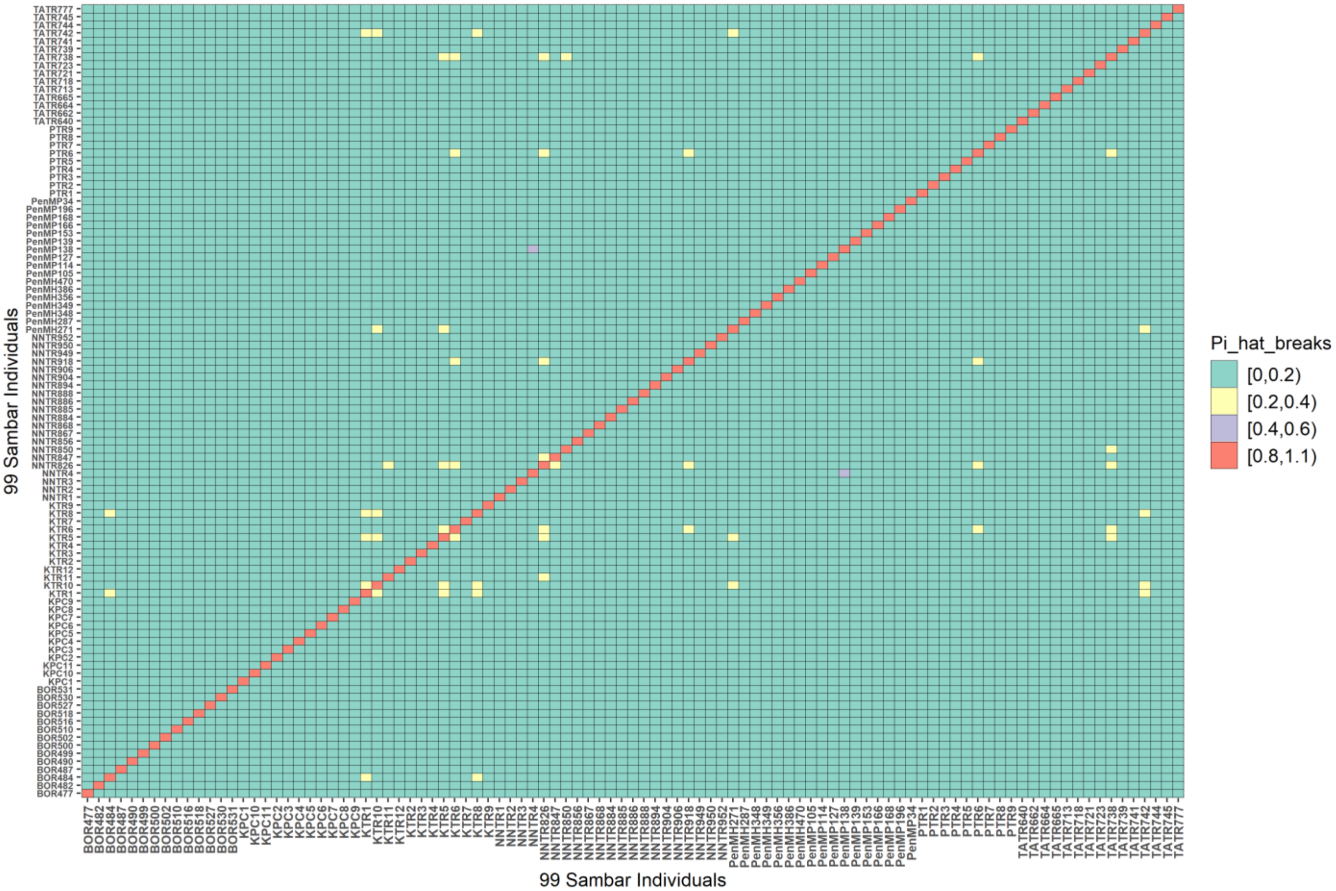

**Figure S5: Rmax and x for the best model:** Relationship of landscape features and maximum resistance offered (x and Rmax) for the best multivariate models for Gaur.

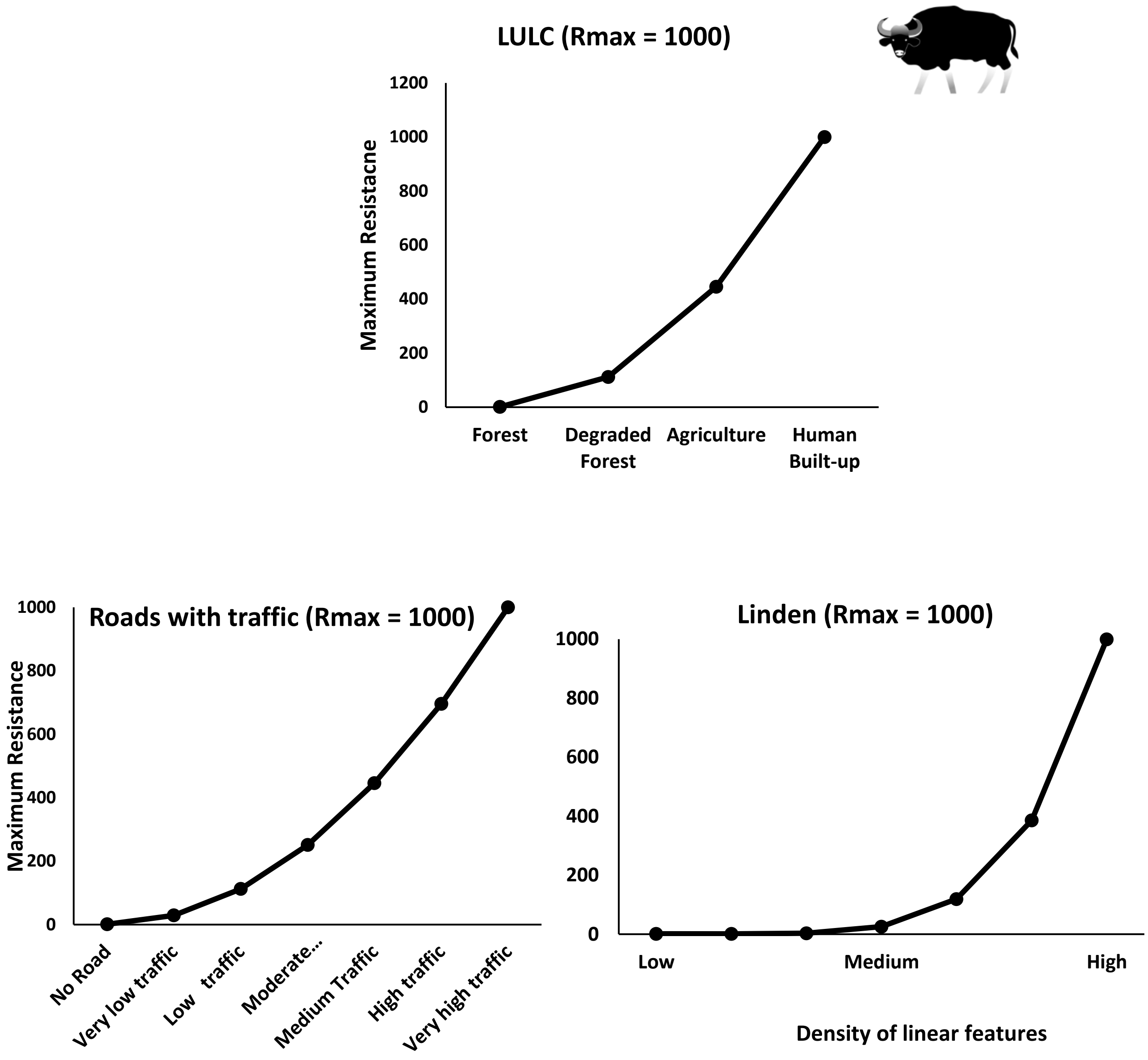

**Figure S6: Rmax and x for the best model:** Relationship of landscape features and maximum resistance offered (x and Rmax) for the best multivariate models for Sambar.

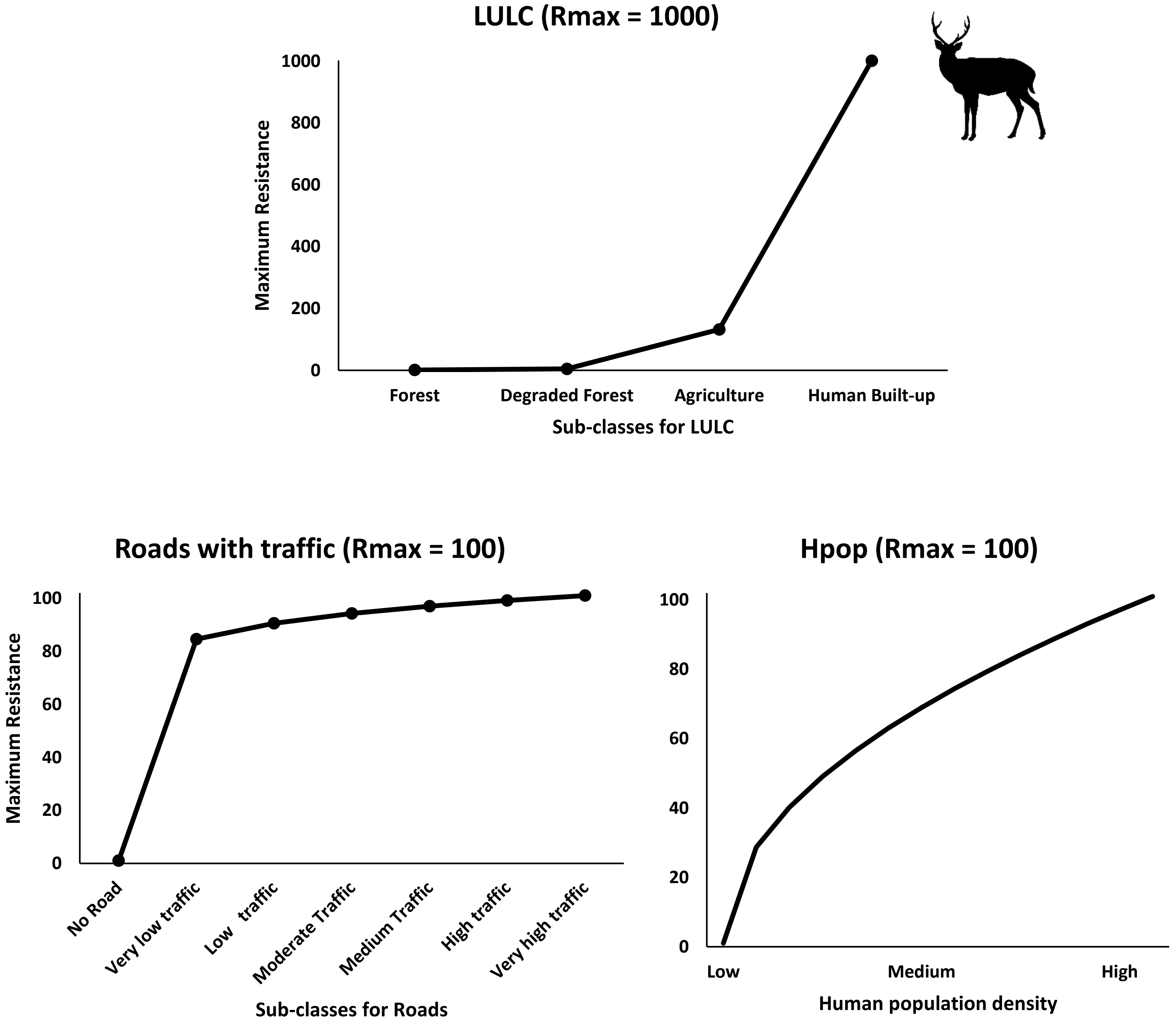

**Figure S7: Plots for estimating the optimal value of K:** The optimum K value for gaur was found to be K=3 and for sambar was K=2. These plots were obtained from a web-based server, CLUMPAK (Kopelman et al., 2015).

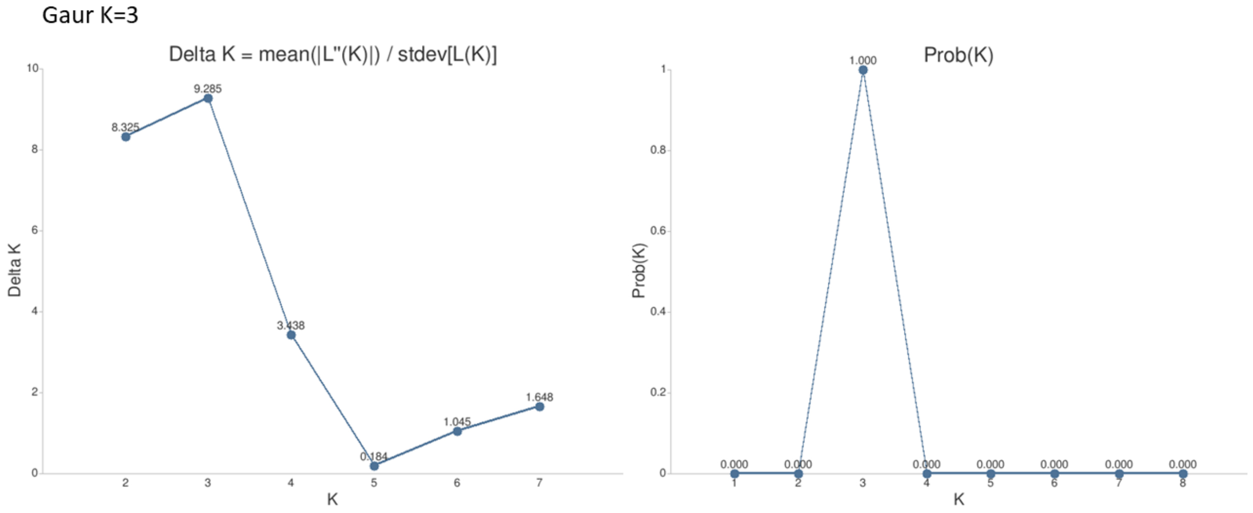

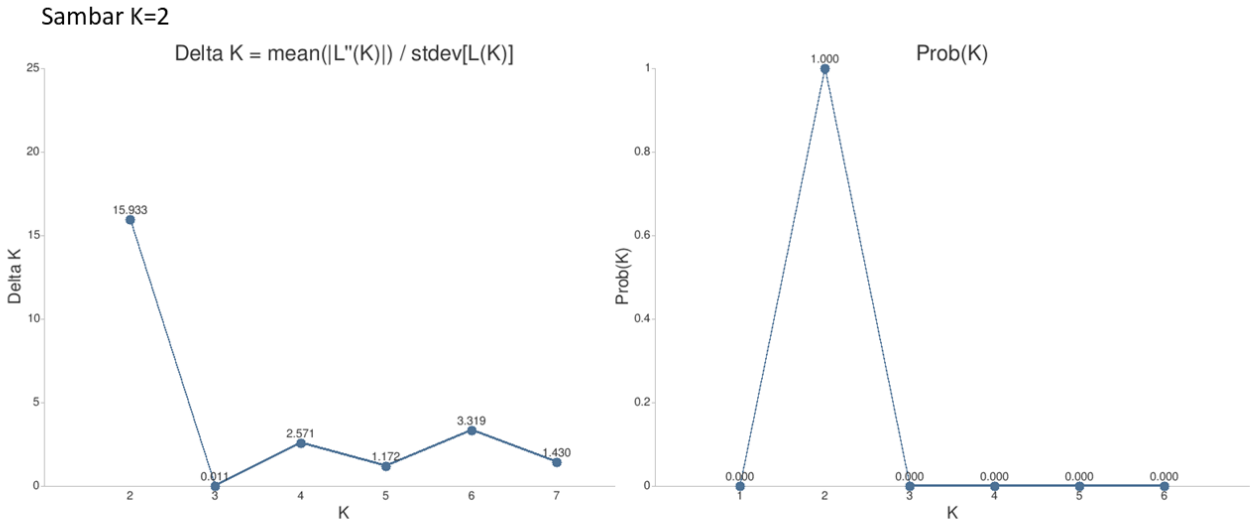

**S2: Detailed ddRAD Library Preparation Protocol**

***Part I. Sample Preparation (Extraction, enrichment and quantification)***

1. DNA Extraction: Extract genomic DNA from fecal samples collected in Longmaire's buffer (Longmire et al., 1997), using Qiagen blood and tissue DNA extraction kit (Qiagen, Toronto, ON, Canada). Use 200 µL of AE buffer for final elusion.
2. Quantify total DNA using Qubit ds-HS kit (Invitrogen, Eugene, OR, USA) and quantify host DNA (hDNA) using qPCR using primers designed for *c-myc* region (Morin et al., 2001).
3. To maximise the hDNA capture, concentrate the samples to the final volume of 20 µL using Cenrivap DNA vacuum concentrator (Labconco).
4. Perform hDNA enrichment following the protocol described in Tyagi et al., 2022 & Chieu and Bergey 2019.
5. Post enrichment, quantify the total and hDNA concentration of the enriched product. To maximise the input DNA amount, concentrate the enriched product volume of 10 µL using Cenrivap DNA vacuum concentrator (Labconco).

***Part II. Reagents preparation (Adapters & Indexes)***

**A. Prepare Adapters** (sequences can be found in Table S3)

1. Make 10X Annealing Buffer stock (adapted from Peterson et al., 2014):
To make 10 mL of 10X annealing buffer use the following reagents in given volumes. For making higher volumes, please modify the proportions accordingly.

| S.No. | Reagent | Volume used | Final conc. |
|---|---|---|---|
| 1 | 1 M Tris-HCl, pH 8 | 1 ml | 100 mM |
| 2 | 5 M NaCl | 1 ml | 500 mM |
| 3 | 0.5 M EDTA | 0.2 mL | 10 mM |
| 4 | $H_2O$ | 7.8 mL | |

Dilute stock to 1X for further use.

2. Resuspend adapters to 100 µM in 10 mM Tris-HCl pH 8 or 1X TE pH 8.
3. To make 100 µL of 40 µM double stranded adapter add:

   40 µL of SphI_Adapter_1A (100 µM)
   40 µL of SphI_Adapter_1B (100 µM)
   10 µL of 10X Annealing buffer (made in step 1)
   10 µL of $H_2O$
   (Repeat the same with MuCI_Adapter_1A and MluCI_Adapter_2A)

4. Incubate the mix at 97.5 °C for 2.5 min, and then cool at a rate not greater than 3 °C per minute until the solution reaches a temperature of 21 °C. Hold at 4 °C.
5. Prepare working strength concentrations of annealed adapters from this annealed stock. The following calculation is for 100 reactions and can be modified accordingly if making for higher and lower reactions.
   a. Make working concentration of Adapters by adding:
      - MluCI Adapter 1 (10 µM): 37.5 µL of 40 µM double stranded MluCI Adapter stock, 112.5 µL 10 mM Tris-HCl.
      - SphI Adapter 2 (4 µM): 5 µL of 40 µM double stranded adapter stock, 45 µL 10 mM Tris-HCl.
      - SphI Adapter 2 (0.1 µM): 2.5 µL of 4 µM SphI Adapter 2, 97.5 µL 10 mM Tris-HCl.

b. Prepare Adapter mix: 5 µL of Adapter Mix is required per sample. For 100 reactions combine:

100 µL of 0.1 µM SphI Adapter 2 (0.02 µM final conc.)
150 µL of 10 µM MluCI Adapter 1 (3 µM final conc.)
250 µL of 1X Annealing Buffer

(***Note:*** Store Adapters and Adapter Mix at −20 °C)

**B. Prepare Indexes** (sequences can be found in Table S4). The indexes can be resuspended in 10 mM Tris-HCl (pH 8) and the main stock should be at a concentration of 100 µM . Make the working solution of 10 µM for indexing PCR (see details below).

***Part III. Library construction (Digestion, Adapter ligation, Indexing & size selection)***

**1. Restriction digestion**

- Make a restriction master mix. (The following calculation is for 1 reaction)

| | | |
|---|---|---|
| 2.0 µL | 10X NEB Cut Smart Buffer | 1X final conc. |
| 0.4 µL | SphI-HF 20U/ µL | 8U |
| 0.4 µL | MluCI-HF 20U/ µL | 8U |
| 7.2 µL | $H_2O$ | |

- Add 10 µL DNA (prepared in Part I) to 10 µL Restriction Master Mix (20 µL total volume).
- Vortex the plate/tubes gently and spin it down
- In a thermocycler, digest for 3 h at 37 °C followed by 20 mins at 65 °C. Hold at 4 °C. Digested products can be stored at 4 °C overnight.

**2. Adapter ligation**

- Make a ligation master mix. (The following calculation is for 1 reaction)

  4.0 µL 10X T4 DNA ligase Buffer

  0.5 µL T4 DNA ligase

  10.5 µL $H_2O$

  5 µL Adapter mix (prepared in Part I: A5b)

- Add 20 µL ligation master mix to 20 µL of digested product.
- Pipette mix the reaction mixture and spin it down briefly.
- Incubate at 23 °C for 2 h in a thermocycler, followed by heat-kill at 65 °C for 10 min.

(***Note***: Move directly to clean-up using AmpureXP beads)

3. **Clean-up of the ligated product:** Using Agencourt AMPure XP Beads (Beckman Coulter, Mississauga, ON, Canada) following the standard protocol (described below) using a 1.5X ratio of bead solution volume to ligation solution volume to remove short DNA fragments such as un-ligated adapters and adapter-adapter ligation products.
   - Bring AMPure XP Beads to room temperature.
   - Vortex the Agencourt AMPure XP bottle for 30 seconds to resuspend magnetic particles.
   - Prepare fresh 80% ethanol (400 µL per sample is required if doing in a 96-well plate).
   - Add Agencourt AMPure XP beads equal to 1.5X the volume of the solution (60 µL of the AMPure beads to the 40 µL of ligated product).
   - Mix thoroughly by pipette mixing 10 times (DO NOT VORTEX).
   - Incubate for 5 min at room temperature.
   - Place the plate onto a magnetic stand for 2 min, or until the solution has cleared.
   - With the plate on the magnetic stand, remove the solution from the reaction plate and discard it. Do not disturb the pellet of separated magnetic beads. If beads are drawn out, leave a few microlitres of supernatant behind.

- With the plate on the magnetic stand, add 200 µL of freshly prepared 80% ethanol to each well of the reaction plate.
- Incubate for 30 s at room temperature. Remove and discard the supernatant.
- Repeat for a total of two washes.
- Be sure to remove all of the residual ethanol from the bottom of the well as it is a known PCR inhibitor and impacts the downstream processing.
- With the plate on the magnetic stand, dry for ~ 2 minutes to ensure all traces of ethanol are removed. (***Note***: Make sure the pallet has not completely dried, it will reduce the elusion efficiency)
- Remove from the magnetic rack and add 20 µL of 10 mM Tris-HCl; pH 8, and pipette mix 10 times.
- Incubate at room temperature for 2 minutes.
- Place the plate onto the magnetic stand for 2 min, or until the supernatant has cleared to separate beads from the solution.
- Transfer the eluent to a new plate or tubes (0.2 mL strip tubes or 96-well plate).

**4. Indexing and PCR amplification**

Before setting up indexing PCR, prepare a sample sheet to plan which indexed primers will be used for each sample (sequence of indexing primers can be found in table S4). (***Note:*** Quantify a few samples before doing the PCR using Qubit DNA HS reagent kit) Prepare the reaction mixture using the following calculations (for 1 reaction):

| | |
|---|---|
| 20 µL | 2X NEB master mix |
| 2.5 µL | Forward primer (i5) |
| 2.5 µL | Reverse primer (i7) |
| 15 µL | Cleaned ligated product |

- The total reaction volume will be 40 µL.

- Pipette mix the reaction using filtered tips and briefly spin it down.
- Split the reaction volume into four aliquots of 10 µL each in different tubes/plates. This is to reduce PCR bias and use these four aliquoted sets for performing PCR.

- Carry out the following PCR program:
  98 °C for 30s
  (98 °C for 30s, 65 °C for 30s, 72 °C for 30s) × 16 cycles
  72 °C for 5 min; Hold at 4 °C

Pool all four aliquots back together and quantify PCR reactions using Qubit DNA HS reagent kit. Compare the concentration of a few samples pre- and post-indexing PCR to check if PCR/indexing has worked.

**5. Dual size selection using AMPureXP beads**

Before adding AMPure XP beads, add 10 µL of $H_2O$ to the indexed products to make the final volume up to 50 µL. To perform a dual-size selection using Agencourt AMPure XP Beads (Beckman Coulter, Mississauga, ON, Canada), the steps are as follows:

- Bring AMPure XP Beads to room temperature and vortex the Agencourt AMPure XP beads for 30 s to resuspend magnetic particles.
- Add 25 µL (0.5X) Agencourt AMPure XP beads and mix thoroughly by pipette mixing 10 times.
- Incubate for 5 min at room temperature.
- Place the plate onto a magnetic stand for 2 min, or until the solution has cleared.
- With the plate on the magnetic stand, transfer the clear solution from the reaction plate to another plate. Discard the beads.
- To this supernatant, add 25 µL of Agencourt AMPure XP beads, making the volume ~100 µL. Mix thoroughly by pipette mixing around 10 times and incubate for 5 min at room temperature.

- Place the plate onto a magnetic stand for 2 min, or until the solution has cleared.
- With the plate on the magnetic stand, remove the solution from the reaction plate and discard it. Do not disturb the pellet of separated magnetic beads. If beads are drawn out, leave a few microliters of supernatant behind.
- For washing of beads with 80% ethanol and elusion follow the procedure described above (Part III: Clean-up)
- For elusion use 20 µL of 10 mM Tris-HCl; pH 8.
- Quantify the size selected libraries using the Qubit DNA HS reagent kit.
- Check the size distribution on a Tapestation 4200 using HS DNA 1000 screen tape to verify the size.

**6. Library normalization and pooling (**adapted from Naik et al., 2023**)**

Calculate individual library concentration in nM, based on the average size and concentration of the library using the Illumina pooling calculator (**https://support.illumina.com/help/pooling-calculator/pooling-calculator.htm**) or using the following formula:

$$\frac{\text{(Concentration in ng/µL)}}{\text{(660 g/mol × average library size in bp)}} \text{ X } 10^6 = \text{concentration in nM}$$

Normalize the concentration of all libraries to 2nM using 10mM Tris-HCl pH 8.0. Pool 2nM normalized libraries by combining 5 or 10 µL of each in one tube. Check the concentration of the pooled library and calculate the nM considering the average size of all libraries. It should be ~2nM.

***Part IV. Prepare final pooled library for sequencing run set-up on NovaSeq 6000***

- Denature and dilute the final pooled library as recommended by Illumina in this guide ***(https://sapac.support.illumina.com/downloads/novaseq-6000-denature-dilute-guide.html).***
- The sequencing run must be set up with the *modified read 1_SPHI* and *read 2_Mluc1* sequencing primer (Table S2). We recommend following the Novaseq custom primer guide ***(https://sapac.support.illumina.com/downloads/novaseq-custom-primers-guide-1000000022266.html)*** for setting up the run.
- Libraries generated using this method cannot be pooled with any other libraries for the sequencing.
